# Amyloid Beta Glycation Induces Neuronal Mitochondrial Dysfunction and Alzheimer’s Pathogenesis via VDAC1-Dependent mtDNA Efflux

**DOI:** 10.1101/2024.05.14.594173

**Authors:** Firoz Akhter, Asma Akhter, Xiongwei Zhu, Hillary Schiff, Arianna Maffei, Justin Douglas, Qifa Zhou, Zhen Zhao, Donghui Zhu

**Author notes:** These authors contributed equally: Firoz Akhter, Asma Akhter. Correspondence (D.Z).

## Abstract

Glycation, the non-enzymatic attachment of reactive dicarbonyls to proteins, lipids, or nucleic acids, contributes to the formation of advanced glycation end-products (AGEs). In Alzheimer’s disease (AD), amyloid-beta (Aβ) undergoes post-translational glycation to produce glycated Aβ (gAβ), yet its pathological role remains poorly understood. Here, we demonstrate that gAβ promotes neuronal mitochondrial DNA (mtDNA) efflux via a VDAC1-dependent mechanism, activating the innate immune cGAS-STING pathway. Using aged AD mice and human AD brain samples, we observed cGAS-mtDNA binding and cGAS-STING activation in the neuronal cytoplasm. Knockdown of RAGE, cGAS, or STING, as well as pharmacological inhibition of VDAC1, protected APP mice from mitochondrial dysfunction and Alzheimer’s-like pathology. Neuron-specific cGAS knockdown confirmed its pivotal role in driving neuroinflammation and cognitive deficits. Treatment with ALT-711, an AGE cross-link breaker, alleviated gAβ-associated pathology. Furthermore, RAGE inhibition in APP knock-in mice suppressed innate immune activation and disease-associated gene expression, as revealed by spatially resolved transcriptomics. Collectively, our findings establish a mechanistic link between gAβ and innate immune activation, identifying VDAC1, the AGE-RAGE axis, and the cGAS-STING pathway as promising therapeutic targets in AD.

**Significance Statement:** This study reveals how a modified form of amyloid-beta disrupts mitochondrial function in neurons, triggering innate immunity and disease progression. We show that this modified amyloid-beta damages mitochondria, activating a specific immune response in the brain. By identifying the key molecules involved, we provide potential targets for new Alzheimer’s treatments aimed at preventing mitochondrial damage and cognitive decline. This research offers fresh insights into Alzheimer’s development and highlights new therapeutic pathways.

## Introduction

Post-translational protein glycation is a non-enzymatic process in which sugars react with proteins, forming molecules known as advanced glycation end-products (AGEs), and it has long-been considered a toxic consequence of carbohydrate metabolism and mitochondrial perturbation(1). This process is implicated in various diseases, including neurodegenerative disorders like Alzheimer’s disease (AD)(2). Both amyloid beta (Aβ) and tau proteins are subject to glycation. Glycated tau in AD induces neuronal oxidative stress and neurodegeneration(3), while the effect of Aβ glycation is underexplored.

Mounting evidence shows Aβ is highly glycated in patients, and senile plaques in AD brains are enriched with AGEs and have nearly three times more AGE adducts than age-matched controls, pointing to a potential involvement in AD, implying that Aβ is glycated(4, 5). Long-lived proteins are favored for AGE production, and Aβ’s stability makes it a suitable substrate for glycation and Aβ-AGE (gAβ) formation(6). Direct covalent crosslinking of gAβ to substrates or its binding to surface receptors, such as receptor for AGE (RAGE), leads to neuroinflammation and synaptic toxicity(7). gAβ with the altered secondary structure is a more suitable ligand than authentic Aβ for RAGE, resulting in the subsequent activation of signaling pathways that can lead to AD-like pathology. RAGE is a known multiligand receptor that can amplify mitochondrial dysfunction(8), innate immune and inflammatory responses as a progression factor(9). Thus, we speculate that gAβ is more prone to bind to/activate RAGE and the downstream innate immune pathways that contribute to AD pathogenesis.

The cGAS-STING pathway is crucial for sensing cytoplasmic dsDNA, triggering the innate immune response(10). Damage-associated molecular patterns (DAMPs), like mitochondrial DNA (mtDNA), can induce inflammation in diseases like AD(11) when mtDNA escapes mitochondria due to stress(12). cGAS, a DNA sensor, activates when mtDNA enters the cytoplasm, leading to inflammasome activation, TBK1 activation, and IRF3 phosphorylation, which contribute to cellular inflammation(13). While STING is highly expressed in microglia, it is also present in neurons(14), though its role in AD remains unclear.

As the most prevalent protein in mitochondrial outer membrane, voltage-dependent anion channel (VDAC) controls cell death, Ca^2+^ influx, metabolism, inflammasome activation, and other processes(15). It has been reported that VDAC oligomer pores allow stressed mitochondria to release mtDNA(16) and Cytochrome C (Cyt C)(17). Cyto C release is a well-established apoptotic event that can lead to caspase 3/7 activation. However, cytosolic Cyto C is also involved in essential cellular processes, such as differentiation, suggesting that its release does not always follow an all-or-nothing pattern and that mitochondrial outer membrane permeabilization does not invariably lead to cell death.(18) Other factors such as inhibitory mechanisms, compensatory responses, or dysfunction of the apoptotic machinery could prevent apoptosis despite increased cytochrome c levels(19, 20). Despite this finding and those in the earlier reports, the notion that VDAC1 might directly mediate mtDNA efflux remains largely unclear. This research explores gAβ’s involvement in inducing neuronal mtDNA efflux dependent on VDAC1 and activating the cytosolic cGAS-STING sensing pathway as well as the subsequent AD pathogenesis.

## Results

### Aβ glycation in human AD brain is linked to mtDNA leakage and cGAS-STING activation

Aβ are increasingly shown to be highly glycated in patients(21, 22), pointing to a potential involvement for this irreversible, non-enzymatic post-translational alteration in AD pathogenesis. In silico modeling confirmed Aβ40 and Aβ42 stability, showing that methylglyoxal (MGO), a toxic metabolite, interacts with both fibrils, with stronger binding to Aβ42 (−2.6) than Aβ40 (−2.2) **(*SI Appendix*, Fig. S.1A, B)**. The formation of *in vitro* glycated synthetic Aβ42 was further validated using a fluorescence spectrophotometer and immunoblot analysis, and by probing and reprobing with anti-6E10 or anti-AGE antibodies **(*SI Appendix*, Fig. S.1C, D)**. The native form lacked AGE enrichment **(*SI Appendix*, Fig. S.1D)**. Higher gAβ plaques, AGE levels, and Aβ-AGE colocalization were observed in the hippocampus **(Fig. 1A, B)** and cortex **(Fig. 1C, D)** of AD samples. RAGE expression was also elevated in the AD cortex **(Fig. 1E)**.

**Fig. 1.**
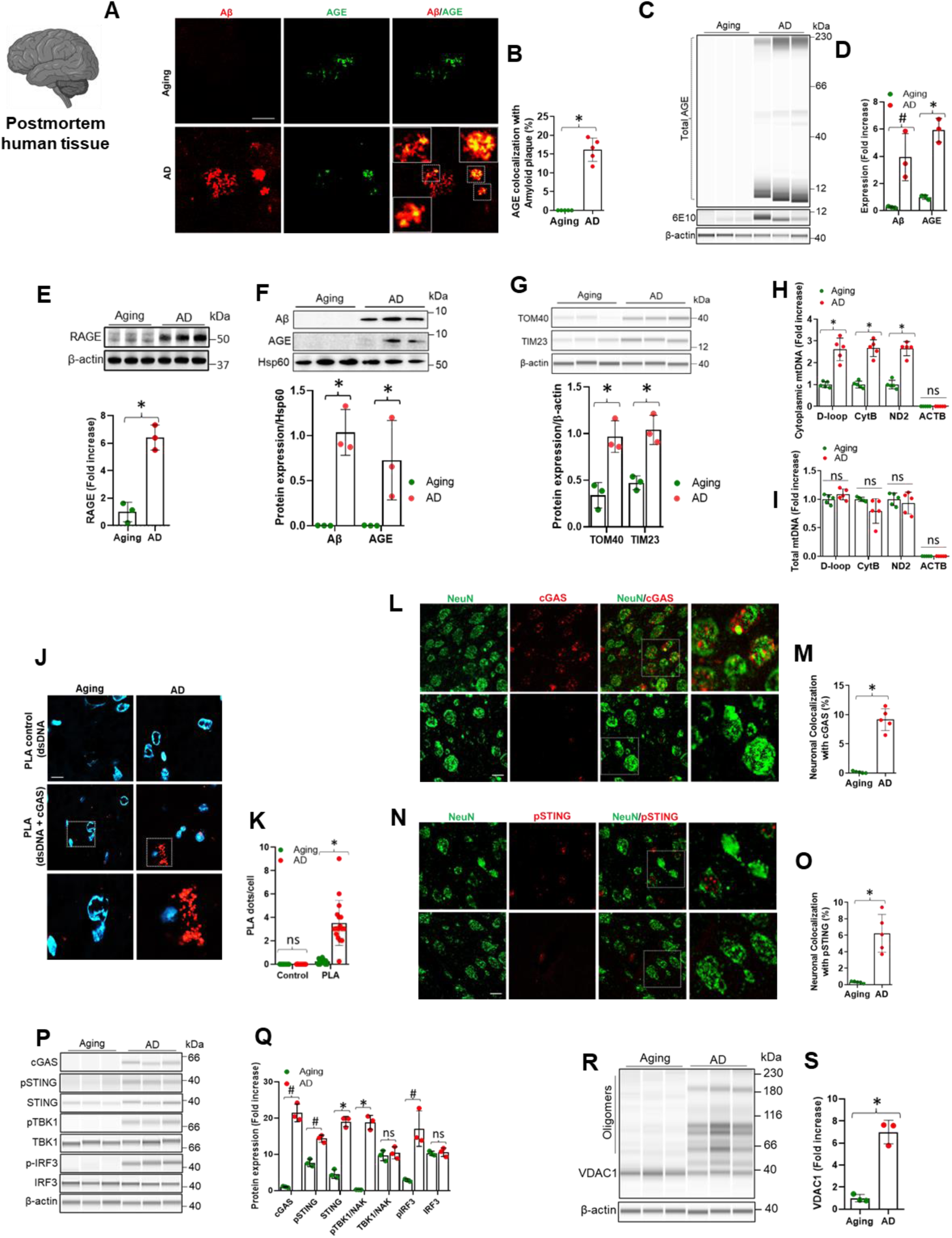
Increased Aβ glycation in human AD brain linked to mtDNA leakage and cGAS-STING activation. (**A**), Immunostaining of Aβ with total AGE in aging or AD human brain hippocampus. Scale bar, 50 μm. (**B**) Percentage AGE colocalization with Amyloid plaque. Mean ± SD; n = 5; *P < 0.001, two-tailed Student’s *t*-test. (**C**), The cortical extracts from aging or AD were immunoprecipitated with AGE and re-probed with Aβ antibody (6E10) (**D**), Quantification of the expression of the total AGE density and Aβ relative to β-actin. Mean ± SD; n = 3; ^#^P < 0.05, *P < 0.01, two-tailed Student’s *t*-test. (**E**), Immunoblotting analysis of the expression of RAGE and quantification relative to β-actin in the human samples. Age-matched normal samples were used as aging controls. Mean ± SD; n = 3; *P < 0.01, two-tailed Student’s *t*-test. (**F**), The purified mitochondria of cortical tissues from aging or AD were immunoprecipitated with Aβ (6E10) and re-probed with AGE antibody and quantification of the expression of the Aβ and AGE relative to Hsp60. Mean ± SD; n = 3; *P < 0.01, two-tailed Student’s *t*-test. (**G**), Immunoblotting analysis of the expression of TOM20 and TIM23 and quantification relative to β-actin in the human samples. Age-matched normal samples were used as aging controls. Mean ± SD; n = 3; *P < 0.01, two-tailed Student’s *t*-test. (**H, I**), Quantification of cytoplasmic and total mitochondrial DNA (mtDNA) by qPCR in indicated AD samples. D-loop, CytB and ND2 are specific fragments or proteins encoded by the mitochondrial genome. ACTB probes for nuclear DNA were used as a control. Mean ± SD; n = 5; ^ns^P > 0.05, *P < 0.01, two-tailed Student’s *t*-test. (**J**), PLA with anti-dsDNA and anti-cGAS antibodies showing the association of cGAS protein with cytosolic or nuclear dsDNA in aging or AD human brain hippocampus. Anti-dsDNA antibody alone was used as the PLA control. Scale bar, 10 μm. (**K**), Quantification of PLA red dots per cell. Mean ± SD; n = 5; ^ns^P > 0.05, *P < 0.01, two-tailed Student’s *t*-test. (**L**), Immunostaining cGAS with NeuN in aging or AD human brain hippocampus. Scale bar, 10 μm. (**M)** Percentage neuronal colocalization with cGAS. Mean ± SD; n = 5; *P < 0.01, two-tailed Student’s *t*-test. (**N**), Immunostaining of pSTING with NeuN in aging or AD human brain hippocampus. Scale bar, 10 μm. Representative images as shown were from the results of five samples in each group. (**O)** Percentage neuronal colocalization with pSTING. Mean ± SD; n = 5; *P < 0.01, two-tailed Student’s *t*-test. (**P, Q**), Immunoblot analysis for the expression and quantification of the cGAS, p-STING, STING, p-TBK1, TBK1, and p-IRF3, IRF3 relative to β-actin. Mean ± SD; n = 3; ^ns^P > 0.05, ^#^P < 0.01, *P < 0.01, two-tailed Student’s *t*-test. (**R, S**), Immunoblotting analysis of the expression and quantification of VDAC1 relative to β-actin in indicated samples of aging or AD. Mean ± SD; n = 3; *P< 0.001, two-tailed Student’s t-test.

It has been shown that Aβ is transported into mitochondria via the translocase of the outer membrane (TOM) and inner membrane (TIM) machinery(23, 24). In AD cortical mitochondrial fractions, gAβ levels (Aβ and AGE co-localization) were higher than controls, with increased TOM40 and TIM23 expression, suggesting glycated amyloid-beta entry via mitochondrial machinery **(Fig. 1F, G)**. Elevated cytoplasmic mtDNA was observed in AD samples, despite similar total mtDNA levels **(Fig. 1H, I)**. PLA showed an over 8-fold increase in cGAS-dsDNA binding in the AD hippocampus **(Fig. 1J, K)**. Immunostaining data confirmed increased levels of cGAS and phosphorylated STING (pSTING) in neurons of the AD hippocampus **(Fig. 1L-O)**.

More importantly, the innate immune cGAS-STING pathway was significantly enhanced in the cortical region of human AD patients, as shown by biochemical analyses on phosphorylated STING, TBK1 and IRF3 **(Fig. 1P, Q)**. Our data showed significantly higher levels of inflammatory cytokines (TNF-α, IL-1β, IL-6) in AD cortical samples **(*SI Appendix*, Fig. S.1E)**, consistent with cGAS-STING activation and the type I IFN pathway (25). Fission and fusion proteins are vital for mitochondrial health(26). Immunoblotting showed increased Drp1/pDrp1 and decreased Mfn-1/2 in AD, indicating mitochondrial changes, along with elevated ROS, impaired glycolysis, and energy imbalance **(*SI Appendix*, Fig. S.1F-N)**. Next, we prepared cytosolic and mitochondrial fractions from cortical tissues from AD **(*SI Appendix*, Fig. S.1O)** and age-matched controls and found much higher levels of cytochrome c (Cyt c) in the cytosolic fraction in AD brains **(*SI Appendix*, Fig. S.1P, Q)**, while Cyt c level in the mitochondrial fraction was higher in controls **(*SI Appendix*, Fig. S.1P, R)**. More importantly, we confirmed higher oligomeric expression of VDAC1 in AD brains **(Fig. 1R, S)**, linking gAβ load in AD with mitochondrial dysfunction and VDAC1 oligomerization, which may facilitate Cyt c release and contribute to cGAS-STING activation in neurons. CytC release is a well-established apoptotic event that can lead to caspase 3/7 activation. However, we observed no significant changes in cleaved caspase 3 or 7 **(*SI Appendix*, Fig. S.1S-V)**, suggesting that the apoptotic cascade is not the primary driver of cGAS-STING activation.

### Aβ glycation exacerbated neuronal mitochondrial stress and neuroinflammation *in vitro*

We investigated whether pathogenic gAβ (Aβ42-AGE) activates cGAS-STING signaling and induces neuroinflammation by culturing primary hippocampal neurons treated with human Aβ (Aβ42) or gAβ (gAβ42). As previously reported(27) RAGE level was increased after Aβ treatment, but we found that gAβ had a much stronger effect in upregulating RAGE in cultured primary neurons **(*SI Appendix*, Fig. S.2A, B)**. Furthermore, we hypothesize that induced expression of TOM40 and TIM23 in gAβ-treated neurons, compared to Aβ, suggests increased transportation of gAβ from neuronal cells to mitochondria via the TOM-TIM mitochondrial machinery **(Fig. 2A-C)**.

**Fig. 2.**
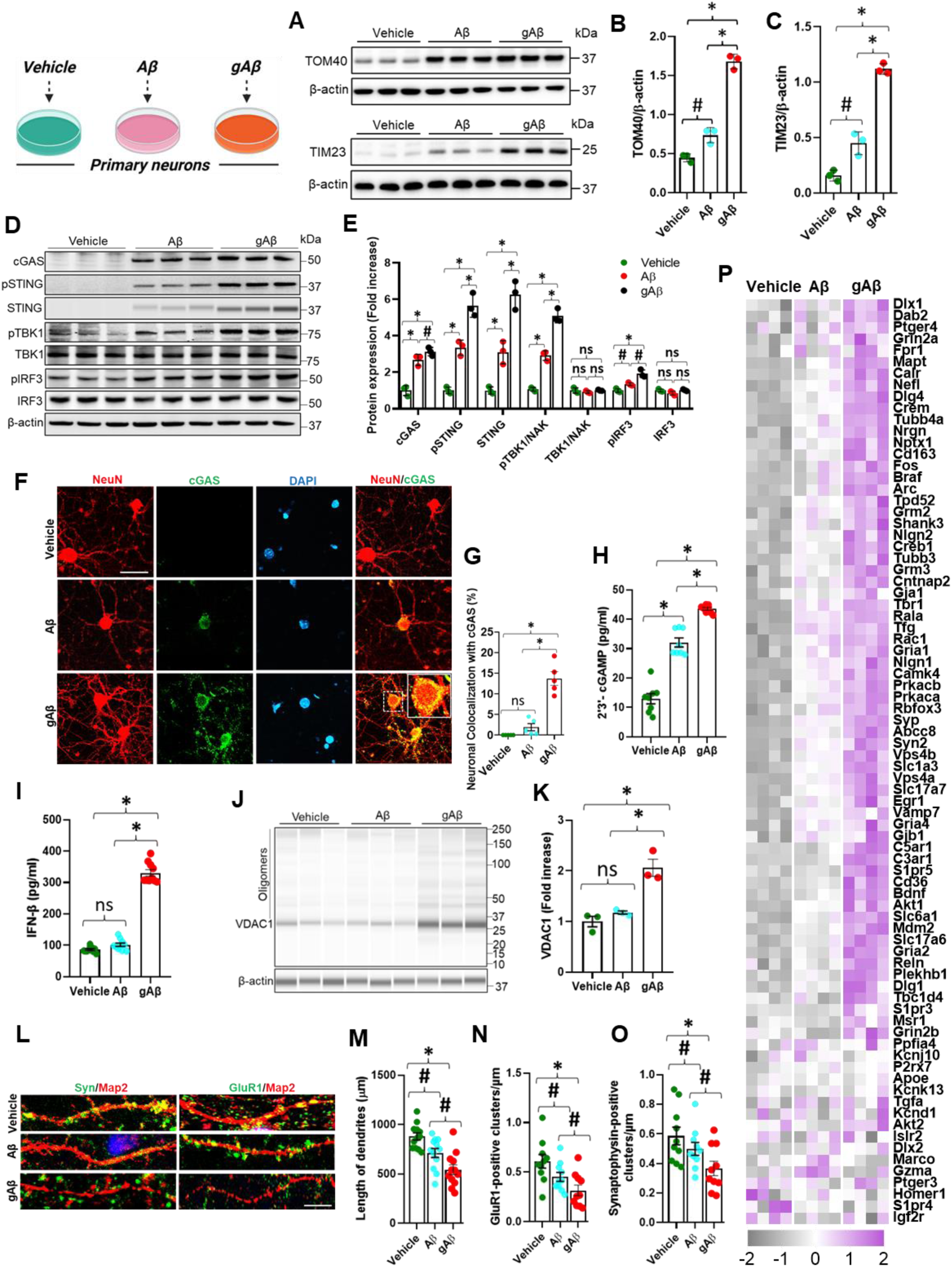
Aβ glycation exacerbates neuronal mitochondrial stress and neuroinflammation *in vitro*. (**A-C**), Immunoblotting analysis of the expression of TOM20 and TIM23 and quantification relative to β-actin in the indicated neurons treated with vehicle, Aβ or gAβ for 24 h. Mean ± SD; n = 3; #P < 0.05, *P < 0.001, one-way ANOVA with Bonferroni’s post hoc test. (**D, E**), Immunoblot analysis for the expression and quantification of the expression of the cGAS, p-STING, STING, p-TBK1, TBK1, and p-IRF3, IRF3 relative to β-actin in the indicated hippocampal neuronal lysate. Mean ± SD; n = 3; ^ns^P > 0.05 ^#^P < 0.05, *P < 0.001, one-way ANOVA with Bonferroni’s post hoc test. (**F**), Immunostaining of cGAS with NeuN in hippocampal neurons (DIV 8), treated with vehicle, Aβ or gAβ for 24 h. Scale bar, 50 μm. (**G**) Percentage neuronal colocalization with cGAS. Mean ± SD; n = 5; ^ns^P > 0.05, *P < 0.001, one-way ANOVA with Bonferroni’s post hoc test. (**H**), ELISA analysis of 2′3′-cGAMP level in vehicle, Aβ or gAβ treated primary neurons with the indicated treatment conditions. Mean ± SD; n = 8; *P < 0.01, one-way ANOVA with Bonferroni’s post hoc test. (**I**), ELISA analysis of IFN-β level in the supernatants of indicated primary neurons treated with vehicle, Aβ or gAβ. Mean ± SD; n = 10; *P < 0.01, one-way ANOVA with Bonferroni’s post hoc test. (**J, K**), Immunoblotting analysis of the expression and quantification of VDAC1 relative to β-actin in indicated neurons (DIV 10) with the same treatment conditions. Mean ± SD; n = 3; ^ns^P > 0.05, *P < 0.01, one-way ANOVA with Bonferroni’s post hoc test. (**L**), Representative images show that gAβ decreased the density of synaptophysin-positive clusters near or on dendrites compared to Aβ or vehicle-treated neurons (DIV 10). Neurons were immunostained with MAP2 and GluR1 as post-synaptic markers, and synaptophysin as a presynaptic marker. (**M-O**), Quantification of Length of dendrites, GluR1-positive clusters and synaptophysin-positive clusters in μm or per μm of dendrite length. Scale bar, 5 μm. Mean ± SD; n = 10; ^#^P < 0.05, \**P* < 0.01, one-way ANOVA with Bonferroni’s post hoc test. (**P**), NanoString nCounter neuroinflammation Profiling panels were employed to analyze neuronal RNA with the indicated treatment conditions. The gene expression heatmap shows selected neuropathology related genes from the NanoString analysis of indicated primary cells (n=4).

Next, we showed that gAβ induced stronger activation of the cGAS-STING pathway, based on the increased levels of cGAS, pSTING, STING, p-TBK1, and p-IRF3 in gAβ-treated neurons relative to Aβ or vehicle **(Fig. 2D, E)**, which was consistent with the fact that the highest cytoplasmic mtDNA level was also seen in gAβ treated neurons **(*SI Appendix*, Fig. S.2C, D)**. The substantial upregulation of cGAS in gAβ treated neurons was further validated by immunocytochemistry **(Fig. 2F, G)**. We also detected increased production of 2′3′-cGAMP by cGAS, in both Aβ and gAβ-treated primary neurons **(Fig. 2H)**, and the level in the gAβ-treated condition was higher than Aβ condition. In examining the consequence of the activation of the cGAS-STING pathway, we found a marked increase in the IFN-β level in the culture media secreted by the neurons treated with gAβ **(Fig. 2I)**, as well as much higher levels of inflammatory cytokines (TNF-α, IL-1β, and IL-6) in gAβ-treated neurons **(*SI Appendix*, Fig. S.2E, F)**. These results suggest that both Aβ and gAβ can induce broad activation of cGAS in neural cells, but gAβ can trigger STING-IFN responses in neurons more robustly.

We further measured that gAβ severely affected mitochondrial dysfunction compared to Aβ-treated neurons **(*SI Appendix*, Fig. S.2G-K)**. Immunoblot analysis revealed increased phosphorylated Drp1 (pDrp1 Se616) and decreased Mfn1/Mfn2, indicating gAβ directly induces mitochondrial dynamics changes **(*SI Appendix*, Fig. S.2L-O)**. Mitochondrial membrane potential, crucial for matrix configuration and Cyt c release during apoptosis(28), was assessed using TMRM and FACS, revealing reduced TMRM levels in gAβ-treated neurons compared to Aβ or vehicle **(*SI Appendix*, Fig. S.3A, B)**. Mitochondria generate ROS, and gAβ-RAGE activation increases H2O2, impairing the electron transport chain(29) (30). gAβ-treated neurons showed significantly higher H2O2 levels than vehicle or Aβ-treated neurons **(*SI Appendix*, Fig. S.3C).** Mitochondrial superoxide is primarily produced in the mitochondria, especially during cellular respiration. Our MitoSOX staining immunofluorescence data demonstrate a significant increase in mitochondrial ROS in gAβ-treated primary neurons compared to vehicle or Aβ treatments **(*SI Appendix*, Fig. S.3D, E)**. FACS analysis and EPR spectroscopy showed elevated ROS in gAβ-treated neurons, with a more pronounced increase compared to Aβ or vehicle **(*SI Appendix*, Fig. S.3F-I)**. Although we observed increased total expression of VDAC1 in glycated Aβ treated neurons **(Fig. 2J)**, VDAC1 oligomerization was not detected **(Fig. 2K)**, suggesting that Aβ may activate the cGAS STING signaling pathway through alternative mitochondrial channels such as the Cyclophilin D containing mitochondrial permeability transition pore, rather than through VDAC1 macropores under acute in vitro conditions.

To analyze whether gAβ and cGAS-STING activation exacerbate neurotoxicity, we cultured mature mouse neurons with vehicle, Aβ or gAβ treatment, and used microtubule-associated protein 2 (MAP2) as a dendritic marker, and synaptophysin and GluR1 as presynaptic and postsynaptic markers, respectively **(Fig. 2L)**. Both Aβ and gAβ treated neuronal dendrites not only showed diminished length of dendrites, but also reduction in GluR1- or synaptophysin-positive clusters **(Fig. 2M-O)**. Remarkably, gAβ had a most deleterious effect on dendrites length and synaptic density compared to vehicles or Aβ treated neurons **(Fig. 2M-O)**. Furthermore, to understand gAβ-induced STING-IFN response in inducing specific neuroinflammation and molecular signatures, we analyzed RNAs from the vehicle, Aβ or gAβ-treated neurons using the amplification-free nCounter neuropathology profiling panel from NanoString. Twenty-nine genes including *Calr*, *Dlg4*, *Egr1*, *Gja1*, *Gria2*, *Mapt*, *Nlgn2*, *Nrgn*, *Plekhb*, *Slc17a7*, *Slc1a3*, *Syn2*, *Syp*, and *Tubb3* out of eighty genes associated with neuropathology were highly expressed in gAβ-treated neurons compared to vehicle or Aβ-treated neurons **(Fig. 2P)**. This data validates that gAβ can cause mitochondria dysfunction and subsequently mtDNA release, and trigger cGAS-STING activation and inflammation in neurons.

### Increased Aβ glycation in APP mice induced mtDNA leakage and cGAS-STING activation

We examined if Aβ glycation worsens mitochondrial dysfunction and AD in vivo. APP mice at 8 months had elevated methylglyoxal (MGO) and Aβ levels compared to controls **(*SI Appendix*, Fig. S.4A-C)**, with increased RAGE expression **(*SI Appendix*, Fig. S.4D)**. Immunofluorescence confirmed Aβ glycation in APP mice **(Fig. 3A, B)**, and gAβ complex analysis showed age-dependent glycation **(*SI Appendix*, Fig. S.4E)**. As mtDNA often binds with TFAM (transcription factor A, mitochondrial)(31), we isolated and characterized nuclear, cytosolic and mitochondrial fractions of brain cells in 8-month-old nonTg and APP mice **(*SI Appendix*, Fig. S.4F)**, and found TFAM proteins were present in the cytosolic fraction from APP mice **(*SI Appendix*, Fig. S.4G, H)**. Furthermore, our immunoblot data not only revealed increased gAβ in the mitochondrial fraction from the cortical brain of APP mice compared to age-matched wild-type controls (nonTg) **(Fig. 3C, D)** but also increased expression of TOM40 and TIM23 **(Fig. 3E, F)**, which may suggest alterations in the mitochondrial import machinery in response to gAβ.

**Fig. 3.**
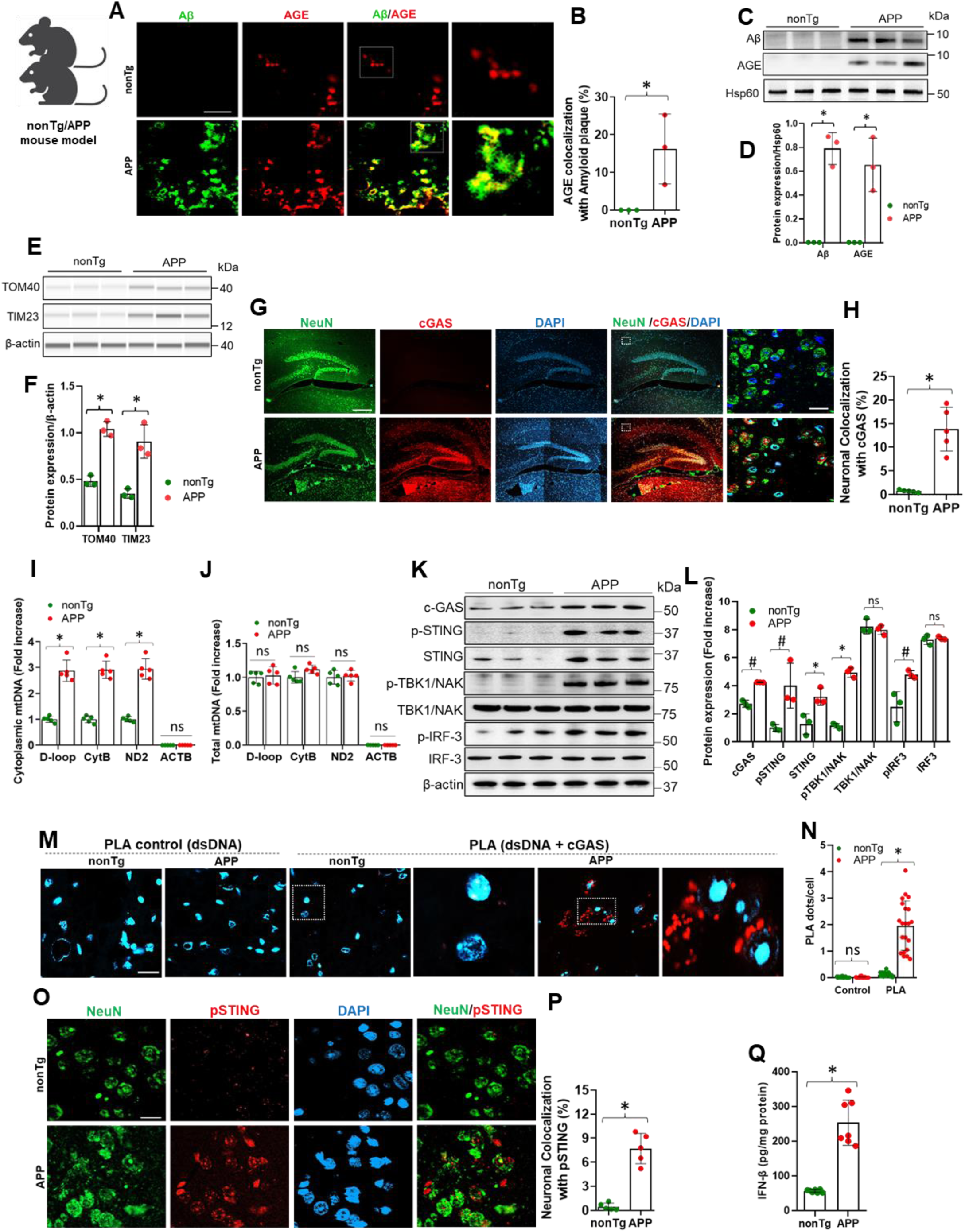
Increased Aβ glycation in APP mice that induces mtDNA leakage and cGAS-STING activation. (**A**), Immunostaining of total AGE with Aβ in 8-month-old nonTg and APP. Scale bar, 100 μm. (**B**) Percentage AGE colocalization with Amyloid plaque. Mean ± SD; n = 3; *P < 0.01, two-tailed Student’s t-test. (**C**), The purified mitochondria of cortical tissues from nonTg and APP mice were immunoprecipitated with Aβ (6E10) and re-probed with AGE antibody. (**D**), Quantification of the expression of the Aβ and AGE relative to Hsp60. Mean ± SD; n = 3; *P < 0.01, two-tailed Student’s t-test. (**E, F**), Immunoblotting analysis of the expression of TOM20 and TIM23 and quantification relative to β-actin in the indicated mice cortical lysate. Mean ± SD; n = 3; *P < 0.01, two-tailed Student’s t-test. (**G**), Immunostaining of cGAS with NeuN in Hippocampus of 8-month-old nonTg and APP mice. Scale bar, 200 μm (low power image) or 10 μm (high power image). (**H**) Percentage neuronal colocalization with cGAS. Mean ± SD; n = 5; *P < 0.01, two-tailed Student’s t-test. (**I, J**), Quantification of cytoplasmic and total mitochondrial DNA (mtDNA) by quantitative PCR (qPCR) in brain cells of 8-month-old APP mice. Age-matched nonTg mice were used as controls. D-loop, CytB and ND2 are specific fragments or proteins encoded by the mitochondrial genome. ACTB probes for nuclear DNA were used as a control. Mean ± SD; n = 5; ^ns^P > 0.05, *P < 0.01, two-tailed Student’s t-test. (**K, L**), Immunoblot analysis for the expression and quantification of the expression of the cGAS, p-STING, STING, p-TBK1, TBK1, and p-IRF3, IRF3 relative to β-actin. Mean ± SD; n = 3; ^ns^P > 0.05, ^#^P < 0.05, *P < 0.01, two-tailed Student’s t-test. (**M**), PLA with anti-dsDNA and anti-cGAS antibodies showing the association of cGAS protein with cytosolic or nuclear dsDNA in hippocampal tissues of 8-month-old APP mice. Anti-dsDNA antibody alone was used as the PLA control. Age-matched nonTg mice were used as control. Scale bar, 10 μm. (**N**) Quantification of PLA red dots per cell. Mean ± SD; n = 5; ^ns^P > 0.05, *P < 0.01, two-tailed Student’s t-test. (**O**), Immunostaining of pSTING with NeuN in Hippocampus of 8-month-old nonTg and APP mice. Scale bar, 10 μm. (**P**) Percentage neuronal colocalization with pSTING. Mean ± SD; n = 5; *P < 0.01, two-tailed Student’s t-test. (**Q**) ELISA analysis of IFN-β level in hippocampus of 8-month-old nonTg and APP mice. Mean ± SD; n = 5; *P < 0.01, two-tailed Student’s t-test.

As mtDNA efflux to the cytosol can be sensed by cGAS and lead to STING activation, we next confirmed that neuronal cGAS expression was significantly increased in APP mice compared to age-matched controls by immunofluorescent staining, indicating an upregulation of the cGAS enzyme in neurons of the APP model (**Fig. 3G, H**). APP mice exhibited markedly higher cytosolic mtDNA levels than nonTg mice **(Fig. 3I, J)**, despite similar total mtDNA. This mtDNA leakage, linked to gAβ and plaque stress, increased with age, as younger (3, 6-month-old) APP mice had lower cytoplasmic mtDNA **(*SI Appendix*, Fig. S.4I, J)**. In 8-month-old APP mice, cGAS, STING, and phosphorylated STING, TBK1, and IRF3 were significantly elevated in the hippocampus compared to nonTg controls **(Fig. 3K, L)**, with weaker activation in younger mice **(*SI Appendix*, Fig. S.4K, L)**. PLA analysis further revealed a >6-fold increase in cGAS-dsDNA interactions in 8-month-old APP mice, most pronounced in the hippocampus and less so in younger cohorts **(Fig. 3M, N; *SI Appendix*, Fig. S.4M, N).** In contrast to their nonTg littermates, APP mice exhibited increased levels of pSTING in their neural cells, suggesting enhanced activation of the cGAS-STING pathway in these neurons **(Fig. 3O, P)**. In addition, the upregulated level of IFN-β was observed in 8-month-old APP mice hippocampus compared to nonTg controls and increased over aging **(Fig. 3Q; *SI Appendix*, Fig. S.4O)**.

We performed bulk gene expression analysis of neuroinflammation in the cortex of 8-month-old AD mice. Compared to nonTg mice, 133 genes showed significant changes: 106 upregulated and 27 downregulated **(*SI Appendix*, Fig. S.4P)**. Pathway analysis revealed neuroinflammatory activities, including cytokine signaling, immune response, apoptosis, and microglial function, suggesting global immune status alterations in the APP model linked to gAβ and cGAS-STING activation **(*SI Appendix*, Fig. S.4Q)**. STING is expressed in the brain with the highest level in microglia, but it also exists in neurons, which is also supported by evidence from iPSC-derived neurons(32).

### Neuron-specific knockdown of cGAS and brain-wide silencing of RAGE, cGAS, or STING attenuate neuroinflammation and AD pathology

To validate the role of cGAS-STING signaling in neurons, we administered either AAV.EB-hSyn-EGFP-m-cGAS-shRNAmir or AAV.EB-hSyn-EGFP-scrmb-shRNAmir intravenously into 6-month-old APP mice to achieve neuron-specific knockdown of cGAS or serve as scrambled controls, respectively **(Fig. 4A)**. Eight weeks post-injection, EGFP expression colocalized with NeuN, confirming successful and selective neuronal transduction **(Fig. 4B)**. Confocal microscopy and PLA staining revealed significantly reduced pSTING expression and decreased dsDNA-cGAS colocalization in NeuN-positive cells within the cortex **(Fig. 4C-F)**. These results were supported by immunoblot analyses showing attenuated activation of the cGAS-STING pathway, including decreased levels of cGAS, pSTING, pTBK1, and pIRF3 **(Fig. 4G, H)**.

**Fig. 4.**
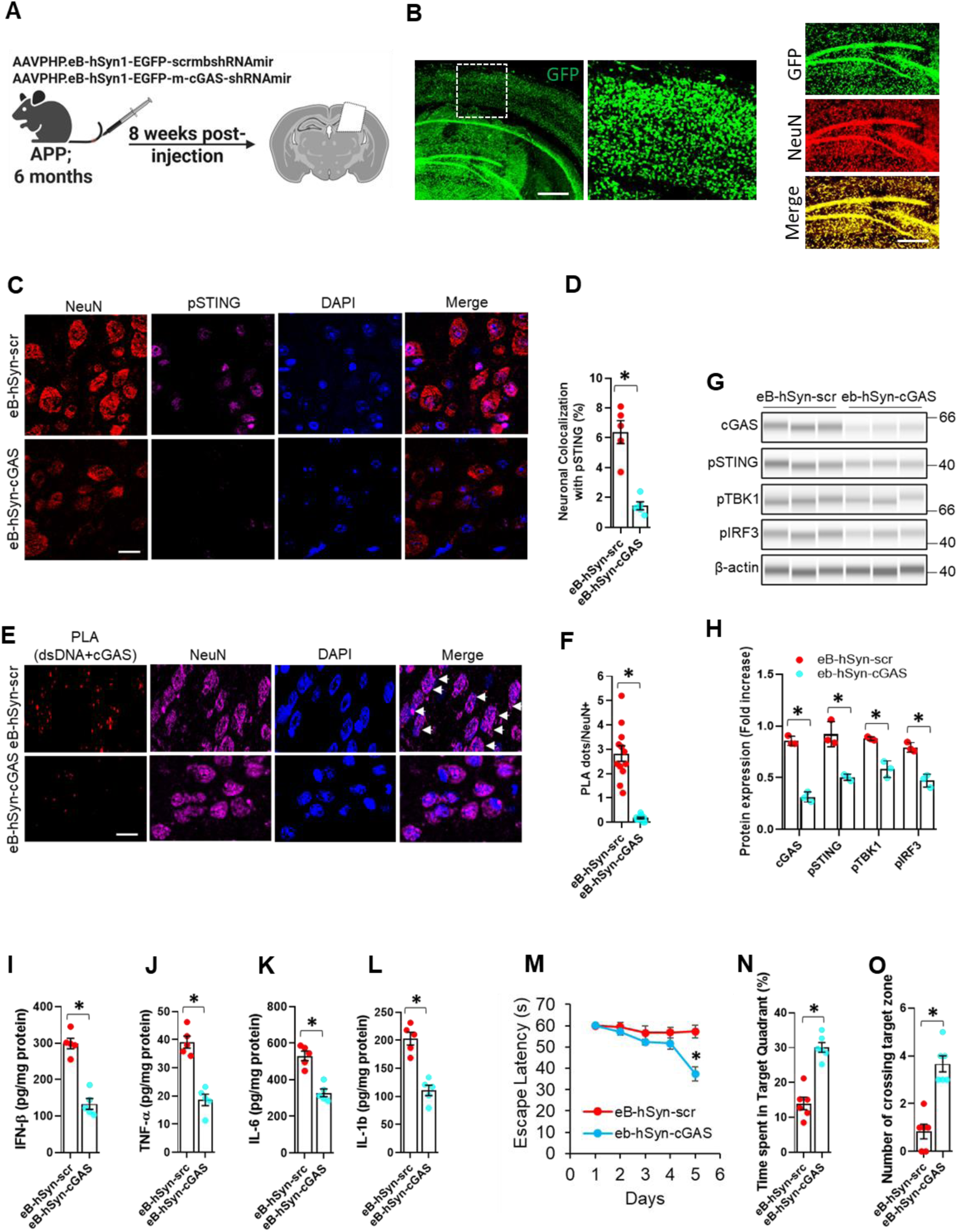
Neuron-specific knockdown of cGAS suppresses cGAS-STING activation, neuroinflammation, and cognitive deficits in APP mice. (A), APP mice were transfected with AAVPHP.eB-hSyn-EGFP-m-cGAS-shRNAmir or scrambled control. (B), Neuron-specific targeting and knockdown were confirmed by GFP and NeuN co-localization. Scale bar, 200 μm (**C**), Immunostaining of pSTING with NeuN in the cortical region of indicated mice. Scale bar, 10 μm. (**D**) Percentage neuronal colocalization with pSTING. Mean ± SD; n = 5; *P < 0.01, two-tailed Student’s t-test. (**E**), PLA with anti-dsDNA and anti-cGAS antibodies with NeuN showing the association of cGAS protein with cytosolic or nuclear dsDNA in neuronal cells in cortical region of the indicated mice. Scale bar, 10 μm. (**F**), Quantification of PLA red dots per NeuN+ cell. Mean ± SD; n = 5; *P < 0.01, two-tailed Student’s t-test. (**G, H**), Immunoblotting analysis of the expression of the cGAS, p-STING, p-TBK1, and p-IRF3 in the indicated samples; n = 3. (**H**), Quantification of the expression of the cGAS, p-STING, p-TBK1, and p-IRF3 relative to β-actin. Mean ± SD; n = 3; *P < 0.001, two-tailed Student’s t-test. (**I-L**), ELISA analysis of IFN-β, TNF-α, IL-6, and IL-1b level in the indicated samples. Mean ± SD; n =5; *P < 0.01, two-tailed Student’s *t*-test. (**M-O**), Escape latency during MWM hidden platform task in indicated mice (M), time spent in the quadrant with the hidden platform (N), mean number of crossings of the target during the probe test (O) of indicated mice Morris Water Maze behavior task of mice. Mean ± SD; n = 6; *P < 0.01, two-tailed Student’s t-test.

Importantly, these molecular changes were associated with a marked reduction in IFN-β expression, as well as other proinflammatory cytokines including TNF-α, IL-6, and IL-1β **(Fig. 4I-L)**, accompanied by improved cognitive performance in the Morris water maze (MWM) testing, indicating behavioral rescue **(Fig. 4M-O)**. Together, these findings demonstrate that neuronal cGAS is a key mediator of gAβ-induced inflammation, mitochondrial dysfunction, and cognitive decline in APP mice.

To further confirm the role of RAGE in gAβ-induced cGAS-STING activation, we administered target-specific shRNA, delivered via adeno-associated viral vector (AAV), intracerebroventricularly (ICV) into 7-month-old APP mice to silence RAGE, cGAS, or STING in the CA1 region of both hippocampi **(*SI Appendix*, Fig. S.5A)**. Immunoblot analysis verified the efficiency of AAV-shRNA delivery and the knockdown of RAGE, cGAS, and STING **(*SI Appendix*, Fig. S.5B-E)**. RAGE knockdown in APP mice restored mitochondrial respiration and ATP production **(*SI Appendix*, Fig. S.5G-H)**, increased DHAP and G3P levels, and significantly reduced intracellular ROS, as indicated by EPR peaks **(*SI Appendix*, Fig. S.5I-L)**. These results indicate that RAGE contributes to mitochondrial dysfunction and cGAS-STING activation. Notably, RAGE knockdown mimicked the effects of cGAS or STING inhibition, suppressing the activation of cGAS, pSTING, STING, pIRF3, and pTBK1 in the hippocampal CA1 region of APP mice treated with AAV-shRNA **(*SI Appendix*, Fig. S.5M-N)**. Knockdown of cGAS also led to a reduction in STING protein levels, suggesting that cGAS activity may influence STING expression in addition to pathway activation(33, 34). Immunofluorescence results further revealed that RAGE and cGAS knockdown significantly reduced cGAS expression in neurons of the hippocampal CA1 region in APP mice **(*SI Appendix*, Fig. S.6A, B)**. This suggests a potential gAβ-RAGE interaction contributes to increased mitochondrial toxicity and immune response. PLA analysis confirmed the substantial reduction of cGAS-dsDNA interaction and the interaction between β III-tubulin (cytoplasmic neuronal marker) and pSTING in the CA1 region of the hippocampus in APP mice following cGAS and RAGE knockdown, or cGAS, RAGE, and STING knockdown **(*SI Appendix*, Fig. S.6C-F)**. Notably, shRNA mediated STING knockdown markedly reduced dsDNA-cGAS PLA signals, suggesting a potential role for STING in regulating cytosolic dsDNA availability or its access to cGAS under stress conditions). Further, we isolated and characterized nuclear, cytosolic, and mitochondrial fractions from the brains of the indicated mice **(*SI Appendix*, Fig. S.6G)**. Immunoblot analyses revealed that mitochondrial stress, indicated by reduced cytosolic cytochrome c and TFAM, was alleviated only in RAGE-knockdown APP mice, suggesting that gAβ-induced mitochondrial dysfunction is RAGE-dependent and occurs upstream of cGAS-STING activation **(*SI Appendix*, Fig. S.6H-K)**, without significant changes in cleaved caspase 3/7 **(*SI Appendix*, Fig. S.6L-N)**.

We used multiplexed gene expression analysis (NanoString nCounter) to examine RAGE, cGAS, and STING knockdown effects in APP mice. Knockdown consistently downregulated genes linked to neuropathology (29 genes, e.g., Akt1, Apoe, Bdnf, Mapt), astrocyte function (18 genes, e.g., C4a, S100b, Slc1a3), microglial function (38 genes, e.g., Il1b, Tnf, Trem2), innate immune response (29 genes, e.g., Ifnar1, Irf3, Tbk1), DNA damage (17 genes, e.g., Bax, Pten), and cellular stress (13 genes, e.g., Atg7, Hdac2) **(*SI Appendix*, Fig. S.7A-F)**. Correlation analysis revealed DNA damage was strongly associated with neuronal and neurotransmission pathways **(*SI Appendix*, Fig. S.7G)**, supporting its role as a driver of neuronal dysfunction in APP mice.

### VDAC1 inhibition restrained mtDNA leakage, cGAS-STING pathway and cognitive decline

Mitochondrial macropore formation, including VDAC oligomers, BAX/BAK macropores, and CypD-containing mPTP(35), facilitates mitochondrial herniation and mtDNA release. To investigate the molecular mechanisms behind mtDNA release and cGAS-STING activation, we treated 6.5-month-old APP mice **(Fig. 5A)** with NSC15364, BIP-V5, or Cyclosporin A (CsA) (36–38) to inhibit VDAC1, BAX, or CypD, respectively. CsA and NSC15364, but not BIP-V5, restored mPTP function, with CsA-treated APP mice showing greater resistance to Ca2+-induced mitochondrial swelling and permeability transition than NSC15364 **(*SI Appendix*, Fig. S.8A, B).** However, CsA did not affect VDAC1 or BAK levels in the APP mice hippocampus **(*SI Appendix*, Fig. S.8C, D)**. Likewise, BIP-V5 treatment suppressed BAX and BAK expression and its oligomers but not CypD in APP mice hippocampus (***SI Appendix*, Fig. S.8E, F**). Our data also showed no changes in Cyt c expression in either characterized fractions or mtDNA-binding protein TFAM expression in the cytosolic fraction after BIP-V5 treatment compared to vehicle-treated APP mice or nonTg control **(*SI Appendix*, Fig. S.8G-K)**. This data suggests that mtDNA efflux is unlikely to be dependent on BAX/BAK/CypD macropores.

**Fig. 5.**
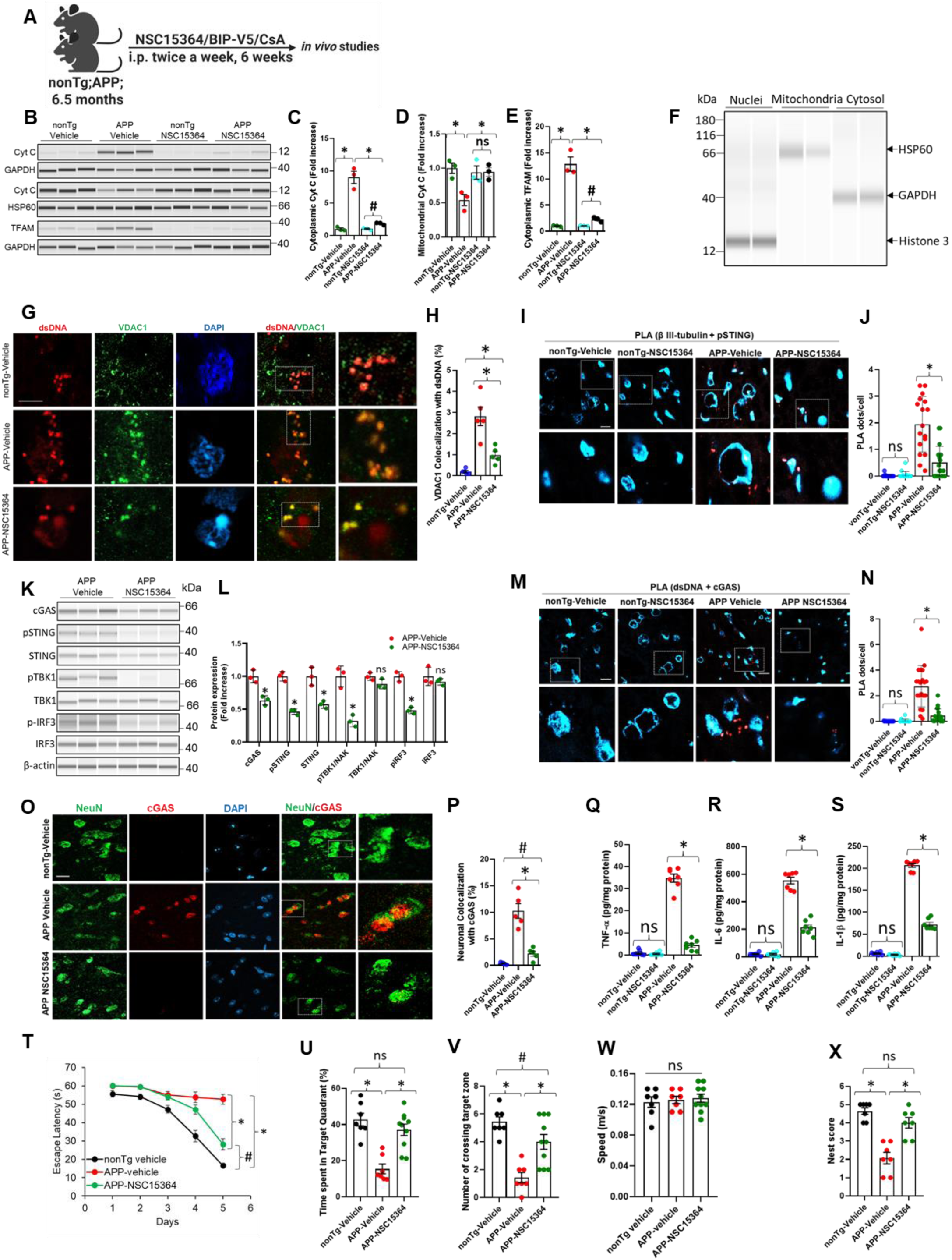
Inhibition of VDAC1 restrains mtDNA leakage, cytochrome c release, cGAS-STING signaling and neuroinflammation in APP mice. (**A**) Schematic representation of BAX, VDAC1, or Cyclophilin D (CypD) inhibitor (BIP-V5, NSC15364, CsA) treatment on 6.5-month-old APP mice. (**B**) Immunoblotting analysis of the expression of the cytochrome c (Cyt c) in the vehicle or NSC15364 administered APP hippocampal cytosolic and mitochondrial fraction and TFAM in cytosolic fraction of indicated samples. (**C, D**), Quantification of the expression of the Cyt c relative to GAPDH or HSP60 in cytosolic and mitochondrial fraction respectively. Mean ± SD; n = 3; ^ns^P > 0.05, ^#^P < 0.05, *P < 0.01, one-way ANOVA with Bonferroni’s post hoc test. (**E**), Quantification of the TFAM relative to GAPDH in cytosolic fraction. Mean ± SD; n = 3; ^#^P < 0.05, *P < 0.01, one-way ANOVA with Bonferroni’s post hoc test. (**F**) Representative immunoblot showing the purity of nuclear, mitochondrial, and cytosolic fractions. (**G**), Immunostaining of dsDNA with VDAC1 in the hippocampal region of indicated mice. Scale bar, 10 μm. (**H**) Percentage VDAC1 colocalization with dsDNA. Mean ± SD; n = 5; ^#^P < 0.05, *P < 0.01, one-way ANOVA with Bonferroni’s post hoc test. (**I**), PLA with anti-NeuN and anti-pSTING antibodies showing the association of neuronal protein with pSTING in hippocampal region of the indicated mice. Scale bar, 10 μm. (**J**), Quantification of PLA red dots per cell. Mean ± SD; n = 5; ^ns^P > 0.05, *P < 0.01, one-way ANOVA with Bonferroni’s post hoc test. (**K**), Immunoblotting analysis of the expression of the cGAS, p-STING, STING, p-TBK1, TBK1, and p-IRF3, IRF3 in the indicated mice hippocampal lysates. (**L**), Quantification of the expression of the cGAS, p-STING, STING, p-TBK1, TBK1, and p-IRF3, IRF3 relative to β-actin. Mean ± SD; n = 3; ^ns^P > 0.05, *P < 0.01, one-way ANOVA with Bonferroni’s post hoc test. (**M**), PLA with anti-dsDNA and anti-cGAS antibodies showing the association of cGAS protein with cytosolic or nuclear dsDNA in hippocampal region of the indicated mice. Scale bar, 10 μm. (**N**), Quantification of PLA red dots per cell. Mean ± SD; n = 3; ^ns^P > 0.05, *P < 0.01, one-way ANOVA with Bonferroni’s post hoc test. (**O**), Immunostaining of cGAS with NeuN in the hippocampal region of indicated mice. Scale bar, 10 μm. (**P**) Percentage neuronal colocalization with cGAS. Mean ± SD; n = 5; ^#^P < 0.05, *P < 0.01, one-way ANOVA with Bonferroni’s post hoc test. (**Q-S**). ELISA for TNF-α (l), IL-6 (m), and IL-1β (n) in indicated mice hippocampal protein. Mean ± SD; n = 7; ^ns^P > 0.05, *P < 0.01, one-way ANOVA with Bonferroni’s post hoc test. (**T-W**), Escape latency during MWM hidden platform task in indicated mice (T), time spent in the quadrant with the hidden platform (U), mean number of crossings of the target during the probe test (V). Average swim speed of nonTg (vehicle-treated) and APP (NSC15364-treated) mice Morris Water Maze behavior task of mice (W). Mean ± SD; n = 10; ^ns^P > 0.05, ^#^P < 0.05, *P < 0.01, one-way ANOVA with Bonferroni’s post hoc test. (**X**) Nest score based on nest building behavior. Nesting deficits indicated by incomplete nests and low nest score. Nesting ability indicated by complete nests, high nest score. Mean ± SD; n = 10; ^ns^P > 0.05, *P < 0.01, one-way ANOVA with Bonferroni’s post hoc test.

Next, we verified that NSC15364, a VDAC1 oligomerization inhibitor, suppressed VDAC1 oligomerization and expression in APP or control mouses hippocampus **(*SI Appendix*, Fig. S.9A, B)**. Interestingly, our data also confirmed the inhibition of Cyt c leakage to cytosolic fraction under the treatment of VDAC1 inhibitor **(Fig. 5B-D)** compared to control, particularly in APP mice. Furthermore, immunoblot data proved a significant reduction of TFAM in characterized cytosolic fraction in VDAC1 inhibitor-treated APP mouse hippocampal cells **(Fig. 5B, E, F)**. Additionally, there was a significant reduction in cytosolic mtDNA in APP mouse brain cells treated with VDAC1 inhibitor, compared to the vehicle treated **(*SI Appendix*, Fig. S.9C, D)**. To confirm whether Cyt C release is linked to the apoptotic cascade, our data show no significant changes in cleaved caspase 3 or 7 **(*SI Appendix*, Fig. S.9E-G)**. This suggests that oligomerization of VDAC1 may facilitate Cyt C release without activating the apoptotic cascade and is not the primary driver of cGAS-STING activation. Furthermore, immunofluorescence revealed colocalization of a flock of dsDNA with VDAC1 clusters in hippocampus of APP mice, which was repressed by the VDAC1 inhibitor NSC15364, validating VDAC1 oligomers dependent mtDNA release **(Fig. 5G, H)**. Additionally, the PLA assay confirmed a significant inhibition of the β III-tubulin and pSTING interaction in the hippocampus of VDAC1 inhibitor-administered APP mice compared to vehicle-treated controls **(Fig. 5I, J)**. This suggests that VDAC1 inhibition may reduce the cGAS-STING pathway activation in neurons, further supporting the role of mitochondrial dysfunction in this process. An Immunoblotting assay showed that cGAS, pSTING, STING, p-TBK1, and p-IRF3 were all significantly reduced in the hippocampal region of 8-month-old APP mice after NSC15364 treatment, compared to vehicle treated groups **(Fig. 5K, L)**. Moreover, PLA assay confirmed a significant inhibition of dsDNA and cGAS interaction in VDAC1 inhibitor administered APP mice hippocampus compared to vehicle treated **(Fig. 5M, N)**. Our immunofluorescence observations further confirmed that the VDAC1 inhibitor NSC15364 blocked cGAS upregulation in hippocampal neurons of APP mice compared to vehicle-treated controls **(Fig. 5O, P)**. Moreover, VDAC1 inhibition by NSC15364 not only effectively inhibited cGAS-STING pathway and production of inflammatory mediators such as TNF-α, IL-6 and IL-1β **(Fig. 5Q-S),** but also the cytosolic release of oxidized mtDNA(39) **(*SI Appendix*, Fig. S.9H)**.

We assessed VDAC1 inhibition in APP mice using the MWM. APP mice showed longer escape latencies and fewer target crossings, indicating memory deficits, which improved with NSC15364 treatment. Swimming speed remained unchanged **(Fig. 5T-W)**. Nest-making ability(40) also improved with NSC15364, indicating enhanced daily task performance **(Fig. 5X)**.

### Pharmacological Disruption of AGE-RAGE Axis attenuates gAβ-Induced VDAC1 Oligomerization, cGAS-STING Activation, and Cognitive Decline in APP Mice

To test whether disrupting the AGE-RAGE axis mitigates gAβ-induced mitochondrial and immune dysfunction, we used ALT-711 (AGE cross-link breaker) and Azeliragon (RAGE antagonist). In wild-type primary cortical neurons, gAβ exposure reduced ATP, increased VDAC1 oligomerization, and activated the cGAS-STING pathway (cGAS, pSTING, STING, pTBK1, pIRF3) with elevated IFN-β. ALT-711 restored ATP production, suppressed VDAC1 oligomerization, and attenuated cGAS-STING activation and IFN-β induction **(*SI Appendix*, Fig. S10A-F)**.

In APP mice treated with ALT-711 for 8 weeks starting at 6 months, brain ATP levels were restored, VDAC1 oligomerization and cytosolic mtDNA were reduced, and cGAS-STING activation and IFN-β levels were suppressed **(*SI Appendix*, Fig. S11A-H)**. ALT-711 treated mice also showed improved nest-building performance **(*SI Appendix*, Fig. S11I)**. These results collectively indicate that ALT-711 mitigates gAβ-induced mitochondrial and immune dysfunction and attenuates AD-like pathology in vivo.

To assess RAGE’s role in cGAS-STING signaling in AD, 6.5-month-old APP mice were treated with the RAGE inhibitor Azeliragon (TTP488) for six weeks **(*SI Appendix*, Fig. S.12A)**. Stereo-seq profiling across cortex, hippocampus, thalamus, striatum, and other regions revealed altered clustering of AD-associated genes in neurons and non-neuronal cells, including markers of mitochondria (VDAC1-3), innate immunity (Irf2, Ifnar1/2, TMEM173), neuropathology (Mapt, Syp, Syn2), A1 astrocytes (C1qa-c, Serping1), disease-associated microglia (Trem2, Apoc1, Cd9), cytokine signaling (Cx3cl1, IL6), DNA damage (BAX, Pttg1), cellular stress (Prdx1, Hdac2), and apoptosis (Bcl2, Map2k4) **(*SI Appendix*, Fig. S.12F-N)**. Immunoblots confirmed suppression of cGAS, STING, and their phosphorylated forms (pSTING, pTBK1, pIRF3) **(*SI Appendix*, Fig. S.13A, B)** and restoration of synaptophysin and PSD-95 **(*SI Appendix*, Fig. S.13C, D)**. PLA assays showed reduced dsDNA–VDAC1 interaction **(*SI Appendix*, Fig. S.13E, F)**, and immunofluorescence confirmed decreased cGAS expression in hippocampal neurons **(*SI Appendix*, Fig. S.13G, H)**. Functionally, Azeliragon improved nest-building **(*SI Appendix*, Fig. S.13I)** and spatial learning and memory in the Morris water maze, with shorter escape latencies and increased target crossings **(*SI Appendix*, Fig. S.13J-L)**, without affecting swim speed **(*SI Appendix*, Fig. S.13M)**. Together, these results suggest that RAGE inhibition mitigates gAβ-induced VDAC1 oligomerization and downstream cGAS-STING mediated neuroinflammation.

## Discussion

Evidence has been presented that Aβ is a stable and suitable substrate for glycation(41). gAβ is one form of AGEs and is more prone to bind to/activate RAGE than authentic Aβ. The plaques in AD brains are highly colocalized with AGEs and are enriched with AGE adducts, implying that Aβ is glycated. However, the role of gAβ in AD pathogenesis was still unclear.

Here, we demonstrate that both MGO-glycated Aβ (gAβ) and authentic Aβ bind to RAGE and can be transported to neuronal cells and further into mitochondria via translocase of the outer membrane (TOM) and inner membrane (TIM). This process induces mitochondrial toxicity, stress, a reduction in mitochondrial membrane potential, and subsequent oligomerization of VDAC1 on the outer mitochondrial membrane. The oligomerization of VDAC1 on the outer mitochondrial membrane can create pores or channels that disrupt normal mitochondrial integrity. This can lead to an imbalance in ion and metabolite transport, causing stress and deformation on the mitochondrial membranes. This stress can result in the protrusion or herniation of the inner mitochondrial membrane through weakened areas of the outer membrane. These changes induce neuronal mtDNA efflux into the cytosol, activating the cytosolic cGAS-STING sensing pathway in AD **(*SI Appendix*, Fig. S.14)**.

Cytosolic mtDNA-cGAS engagement in neurons leads to STING activation and downstream interferon signaling in aged AD mice and human AD brains. Supporting this, both genetic silencing and pharmacological inhibition of the RAGE-cGAS-STING axis (including RAGE, cGAS, STING, or VDAC1) rescued mitochondrial integrity, reduced neuroinflammation, and attenuated AD like pathology. Notably, neuron-specific knockdown of cGAS alone was sufficient to reduce STING pathway activation, neuroinflammation, and restore cognitive function in APP mice. This emphasizes the neuronal origin of innate immune signaling and highlights cGAS as a neuron-intrinsic sensor of mitochondrial stress in AD.

Pharmacological modulation of AGE-RAGE signaling has been previously suggested as a potential therapeutic strategy in age-related diseases, including AD(42, 43). In this context, targeting the AGE-RAGE pathway with ALT-711, a well-characterized AGE cross-link breaker, provided both mechanistic insight and therapeutic promise. In gAβ-exposed primary neurons, ALT-711 restored ATP levels, reduced VDAC1 oligomerization, and suppressed activation of the cGAS-STING pathway and IFN-β induction. When administered to APP mice, ALT-711 similarly decreased cytosolic mtDNA leakage, attenuated cGAS-STING signaling, and improved behavioral performance, demonstrating in vivo rescue of mitochondrial and immune function. These findings underscore the therapeutic potential of targeting AGE-RAGE mediated mitochondrial stress to prevent downstream neuroinflammatory cascades in AD. Furthermore, our spatial transcriptomic profiling following Azeliragon treatment, a small-molecule RAGE antagonist further supports a RAGE dependent mechanism linking gAβ to innate immune activation. Azeliragon reversed AD associated transcriptional signatures across neuronal and non-neuronal populations and reduced cGAS STING pathway activity in the hippocampus. These results, in combination with ALT-711 data, solidify the central role of RAGE in mediating gAβ induced mitochondrial and innate immune perturbations in AD.

Cytosolic mtDNA leakage and cGAS-STING activation in AD has been confirmed in microglia, where STING is highly expressed. Interestingly, we found that STING is also highly expressed in the neurons of AD patients and APP mice but not in normal control counterparts, suggesting neuronal STING is only expressed under stress, in line with earlier reports that STING can also be detected abundantly in neurons under certain conditions. Previous studies found the STING-IFN cascade was present mainly in microglia(10, 44); however, our results raise the question of how the signal of cGAS sensing dsDNA within neural cells is transduced to the activation of the microglial STING-IFN cascade. By RNA-sequencing, we postulated that the activation of the IFN response was only present in gAβ-stressed neurons but not authentic Aβ. It is suggested that directly activating neuronal STING by cGAMP promotes axon regeneration and can be transported between neighboring cells via specific transporters and gap junctions(45, 46). Consistent with this, we proposed that DNA binding cGAS in gAβ-stressed neurons undergoes phase separation and generates the 2’3’-cGAMP, which not only translocate from the ER towards the ER-Golgi intermediate compartment to activate STING in neurons but also astrocytes and microglia and subsequently induces the activation of the STING-IFN cascade, thereby stimulating/amplifying neuroinflammatory responses.

RAGE contributes to AD pathology, in part, by being involved in the translocation of gAβ from the extracellular to the intracellular space and transported into mitochondria via the TOM-TIM mitochondrial machinery, thereby enhancing gAβ cytotoxicity via mitochondrial dysfunction(23). Remarkably, through mitochondrial VDAC1 expression and homo-oligomerization mitochondrial pores, mtDNA is released into the cytosol in response to pathogenic stimuli(47, 48). We found that only gAβ, not authentic Aβ, can induce mtDNA efflux in neurons, which is dependent on VDAC1. The adenine nucleotide translocase, CypD, and the VDAC are hypothesized to be involved in the mPTP in the mitochondrial matrix(49). Consistent with this we observed a correlation in mitochondrial swelling between them, however, mtDNA leakage was independent of CypD. In addition, our data show increased cytosolic cytochrome c in human AD and APP mice samples compared to controls; however, we observed no significant change or compromised activation of caspase-3/7. This indicates that although cytochrome c release occurs as part of mitochondrial stress, it does not necessarily lead to full apoptotic activation. Instead, cytoprotective mechanisms may counteract caspase activation. In this context, the DNA-cGAS-STING pathway may not be a direct secondary response to apoptosis but rather an alternative or compensatory signaling response, potentially triggered by mitochondrial dysfunction and oxidative stress without full progression to apoptosis. We presented the first evidence of VDAC1 directly causing mtDNA efflux in an amyloidogenic mouse model. In addition, we show a connection between gAβ-RAGE and cGAS-STING sensing, and mitochondrial dysfunction. Our work provides a mechanistic description of VDAC1-mediated mtDNA release under gAβ and subsequent mitochondrial stress/apoptosis, which represents one of the innate immune cGAS-STING activations. While this work delineates a key pathological pathway, further investigations are needed to fully elucidate the intercellular transfer of cGAMP and its role in activating the microglial STING-IFN cascade. Future research is necessary to better understand how the neurodegenerative changes are modulated by innate or adaptive immunity and whether these detrimental effects on neuronal function can be partially or even completely reversed.

## Materials and Methods

We used post-mortem human temporal cortex samples categorized by neuropathology and Braak stages, and the AppSAA KI/KI mouse model to assess amyloid pathology and gene expression changes. Primary hippocampal neurons from P0 pups were treated with Aβ or gAβ for in vitro studies. APP mice received intracerebroventricular injections of AAV9-shRNA constructs targeting RAGE, cGAS, and STING, and neuron-specific knockdown of cGAS was achieved using AAV.hSyn-cGAS-shRNA. Additionally, the AGE-RAGE pathway and VDAC1-mediated mitochondrial dysfunction were pharmacologically targeted to evaluate their therapeutic relevance. Techniques included NanoString gene expression analysis, immunofluorescence, immunoblotting, spatial transcriptomics (Stereo-seq), and behavioral assessments (MWM, Nest test), with methodologies detailed in the SI Appendix. ***SI Appendix***.

### Statistics

The sample size was not predetermined using any statistical techniques. Except where noted specifically, neither humans nor animals were excluded from the analysis. All appropriate data were analyzed using StatView statistics software or GraphPad Prism 8. All hypothesis tests were performed as two-tailed tests. Specific statistical analysis methods are described in the related figure legends where results are presented. All data were expressed as the mean ± SD or ± SE. Values were considered statistically significant for P values <0.05.

## Acknowledgements

We thank the patients and families for their tissue donations. We also appreciate the NMR Laboratory at the University of Kansas for EPR assistance and the CMIC and Flow Cytometry Core at Stony Brook University for microscopy and FACS support. Illustrations were created with BioRender.

## Author contributions

D.Z. and F.A. conceptualized the study. F.A. and A.A. performed experiments, analyzed data, and wrote the manuscript. X.Z. provided postmortem human tissue samples from AD patients and controls. H.S. conducted viral knockdown experiments. J.D. performed EPR measurements. X.Z., A.M., Q.Z., Z.Z., and D.Z. provided manuscript feedback. D.Z. secured funding.

