## Supplementary material for "Amyloid Beta Glycation Induces Neuronal Mitochondrial Dysfunction and Alzheimer’s Pathogenesis via VDAC1-Dependent mtDNA Efflux": SI Appendix

**S1 Appendix:**

**Material and Methods.**

**Human tissues.** Post-mortem human brain samples from the temporal cortex (including Brodmann area 38) were obtained from Case Western Reserve University. The postmortem interval (PMI) was < 9 hours. Informed consent was obtained, and the study approved by the Institutional Review Board of Case Western Reserve University, Cleveland, Ohio. Brain samples were categorized based on neuropathological evaluation and Braak stages. Samples from aged subjects without AD or other neurodegenerative diseases served as age-matched controls. Tissues were rapidly frozen for immunoblotting and fixed in paraffin for immunohistochemistry, with 5-μm sections used. All data presented in the figures are derived from human tissue samples representing distinct patients with varying clinical backgrounds, with an average age of 78 ± 8.5 years, including 18 females and 22 males. This ensures that the data reflects diverse clinical conditions and that the findings are not influenced by repeated samples from the same patient. All procedures adhered to Case Western Reserve University’s ethical standards.

**Mice**

Mouse studies were approved by the Animal Care and Use Committee of Stony Brook University in accordance with the US National Institutes of Health guidelines for animal care. The App knock-in mouse model (App^SAA^ KI/KI; B6.Cg-Apptm1.1Dnli/J, Jackson Lab # 034711) of Alzheimer’s disease (AD) was used which carries humanizing Aβ region R684H, F681Y, and G676R mutations, and the KM670/671NL (Swedish) mutation in exon 16 as well as the E693G (Arctic) and T714I (Austrian) mutations in exon 17 of the mouse App (amyloid beta (A4) precursor protein) gene. Mice recapitulate amyloid plaque pathology in vivo without disturbing intrinsic gene expression. Both male and female were used throughout the study. The 7-months old App^SAA^ (APP) mice were used for RAGE-shRNA, cGAS-shRNA, TMEM173-shRNA and scrambled intracerebroventricular injections. The 6-months old APP mice were used for AAVPHB.eB-hSyn1-EGFP-m-cGAS-shRNAmir and scrambled intravenous (i.v.) injection. The 6.5 months-old APP mice were intraperitoneally (i.p.) injected with Bcl-2-associated X protein (BAX), voltage-dependent anion channel (VDAC), Cyclophilin D (CypD) or RAGE inhibitor i.e., BIP-V5 (5 mg kg−1 BW; 10% DMSO + 40% PEG300 + 5% Tween-80 +	45% saline; HY-P0081, MCE), NSC 15364 (2.5 mg kg−1 BW; 10% DMSO + 40% PEG300 + 5% Tween-80 +	45% saline; Cat #HY-108937, MCE), Cyclosporin A (CsA) (1 mg kg−1 BW i.p. 5% DMSO	40% PEG300	5% Tween-80	50% saline; Cat #HY-B0579, MCE) twice each week, Azeliragon (100 mcg; 10% DMSO	40% PEG300	5% Tween-80	45% Saline; Cat #HY-50682, MCE) every day for 6 weeks or ALT-711 (1 mg kg−1 BW; 10% DMSO + 90% corn oil; HY-106024B, MCE) every day for 8 weeks starting at 6 months of age. The solvent of the inhibitor was used as vehicle control with the same volume for i.p. injection for 6.5 or 6-month-old APP mice. Animals were maintained in facilities with a 12-h light–dark cycle, an average ambient temperature of 21°C, an average humidity of 48% and had free access to food and water. The Institutional Animal Care and Use Committee of Stony Brook University approved all experimental procedures and protocols. All guidelines for research with animals were adhered to during this study.

**Primary neuron cell cultures.** Mouse pups (P0) were decapitated, and hippocampi were dissected and treated with 0.25% trypsin. After trituration, cells were plated on poly D-lysine-coated cover slips in high glucose DMEM with 10% fetal calf serum (Invitrogen). Cells were incubated at 37°C with 5% CO2. The following day, the medium was replaced with Neurobasal A supplemented with 2% B27 and 2 mM GlutaMAX (Life Technologies). After 10 days in vitro (DIV), half of the medium was replaced with fresh B27-Neurobasal A containing 2 µM Aβ, 2 µM gAβ, or vehicle for 24 hours.

**Glycated amyloid beta (****gAβ) preparation.** Glycated Aβ (gAβ) was prepared by incubating 10 mg/ml of human Aβ1-42 (Sigma-Aldrich) with or without 0.5 mM methylglyoxal (MGO) in 0.1 M phosphate buffer, pH 7.2, at 37^o^C for three weeks under sterile conditions. After dialysis against phosphate buffer for 48 h at 4^o^C to remove MGO, the prepared glycated Aβ (gAβ) was sterilized by filtration and kept at -20^o^C in aliquots. The production of gAβ was identified with a fluorescence spectrophotometer measuring AGE-specific fluorescence at an emission of 440 nm and excitation of 370 nm (Perkinelmer, Waltham, MA, USA).

**Quantitative real-time PCR.** Total RNA was extracted using the PureLink™ RNA Mini Kit (Thermo Fisher Scientific) and treated with RNase-free DNase (Promega) for purification. RNA quantity and quality were assessed by Nanodrop. cDNA was synthesized from 1 µg of RNA using the M-MLV Reverse Transcriptase kit (Promega). Target gene transcripts were quantified by qPCR with PereCTa SYBR Green SuperMix (Quanta Biosciences) on a CFX96 real-time PCR system (Bio-Rad).

**Quantification of mtDNA release.** Primary neurons (2 × 10^6^) or adult mouse hippocampi were resuspended in digitonin buffer (150 mM NaCl, 50 mM HEPES, pH 7.4, 25 µg/ml digitonin; Millipore Sigma). After incubation and centrifugation, a 1:20 dilution of the supernatant was used for qPCR. The pellet was resuspended in lysis buffer (5 mM EDTA, proteinase K; Qiagen), incubated at 55°C overnight, diluted, and heated at 95°C. Samples were then used for qPCR with mtDNA-specific primers (Dloop1, CytB, Nd2), as listed in ***SI Appendix*, Table S2**

**Immunoblotting.** Protein samples were separated by SDS–PAGE and transferred to a nitrocellulose membrane (Amersham). The membrane was blocked in 5% skim milk in 1× PBST for 1 h and incubated overnight with primary antibody in 5% BSA at 4 °C. The membrane was incubated with anti-mouse or anti-rabbit HRP-conjugated secondary antibodies in 5% BSA and bands were developed with FluorChem imaging system (ProteinSimple). Immunoblot was also run by advanced western blot Abby (ProteinSimple) as per the kit instruction. All antibodies used are listed in ***SI Appendix*, Table S4**.

**Immunofluorescence.** PBS-perfused mice were fixed in 4% PFA overnight, followed by immersion in 30% (w/v) sucrose solution for 48 h and frozen with liquid N2. Samples were stored at −80°C prior to sectioning. The frozen brain was cryostat sectioned (coronal section) into 20 μm thick sections onto slides for histological staining. Sections were blocked for 1.5 h in PBST (containing 2% (w/v) BSA and 0.5% (v/v) Triton-X) followed by 1 h incubation with 0.5% (v/v) Mouse on Mouse (M.O.M.™) Blocking Reagent (Vector Laboratories). The primary antibodies above were diluted in 3% donkey serum in 1× PBST (0.2% Triton-X 100) and incubated overnight at 4°C. Slides were then washed, and immunoreactivity was detected using donkey anti-rabbit IgG Highly Cross-Adsorbed Secondary Antibody, Alexa Fluor Plus 488, donkey anti-Mouse IgG Highly Cross-Adsorbed Secondary Antibody, Alexa Fluor Plus 595 for 1 h at room temperature. The sections were mounted in SlowFade Gold Antifade Reagent with DAPI (Molecular Probes, Thermo-Fischer, Waltham, USA) and sealed by use of nail polish. Slides were visualized using a Zeiss LSM 980 Airyscan 2 NLO Two-Photon Confocal Microscope and then analyzed by ImageJ software. Detailed information of primary and secondary antibodies used is listed in ***SI Appendix*, Table S4**.

**Production of AAV9-RAGE/cGAS/TMEM173 construct.** Viral production and packaging using recombinant adeno-associated virus type 9 (rAAV9) encoding a protein of interest under control of the type III RNA polymerase III (U6) promoter (Vector Biolabs). Briefly, AAV-GFP-U6-shRNA plasmids were co-transfected into HEK293 cells with pAdhelper and AAV9-RC plasmid. 2 days after transfections, cell pellets were harvested, and viruses were released through 3x cycles of freeze/thaw. Viruses were purified through CsCl-gradient ultra-centrifugation, followed by desalting. Viral titer (GC/ml - genome copies/ml) are determined through real-time PCR.

**Short hairpin RNA (shRNA) Intracerebroventricular Injection and cell transfection**

RAGE small hairpin RNA (shRNA) (AAV9-GFP-U6-m-RAGE-shRNA, target sequence: AGCTGGCACTTAGATGGGAAACTTCAAG AGAGTTTCCCATCTAAGTGCCAGCT), cGAS shRNA (AAV9-GFP-U6-m-cGAS-shRNA, target sequence: GAGCTACAAGAATATTATGAAATTCAAGAGATTTCATAATATTCTTGTAGCTC) and STING shRNA (AAV9-GFP-U6-m-TMEM173-shRNA, target sequence: CAACATTCGATTCCGAGATATT TCAAGAG AATATCTCGGAATCGAATGTTG) were used in our experiments. The RAGE, cGAS, STING and scrambled shRNA (AAV-GFP-U6-shRNA) were synthesized by Vector Biolabs (Pennsylvania, USA). Male and female APP mice at 7-month of age were anesthetized with an intraperitoneal (IP) injection of a cocktail containing 100mg/kg ketamine and 10mg/kg xylazine. Bupivacaine (2.5mg/ml, ~0.1mL) was administered under the scalp for local anesthesia. Craniotomies were made over the left and right CA1 at the following coordinates: -2.0mm posterior from Bregma, 1.1mm and 1.5mm lateral from midline (2 separate injection sites/side). A pulled glass pipette was attached to a nanoject pressure injector (Drummond nanoject II) and filled with shRNA virus (AAV9-GFP-U6-m-RAGE-shRNA, AAV9-GFP-U6-m-cGAS-shRNA and AAV9-GFP-U6-m-TMEM173-shRNA). The pipette was lowered 1.2mm below the pia. Approximately 1.0µl total volume was injected per side (0.5µl/injection site) over 10 minutes. The pipette tip was left in place at least 10min after the end of the injection. The skin was then sutured closed, and the animal recovered on a heating pad. Mice were then returned to their home cage and monitored. Vybrant Dil Cell-labeling Solutions (Thermo Fisher) was used to confirm the location of viral delivery.

**Docking study**

The structure of amyloid beta fibril (40 and 42 residues) was obtained from the protein data bank using the PDB IDs 2M4J (40 residues) and 8eze42 (42 residues). The PubChem database was accessed to find the 3D Conformer of Methylglyoxal (MGO). These structures were prepared for docking analysis using Discovery Studio 2021, and the prepared Methylglyoxal was docked with both the structure (40 and 42 residues) of amyloid beta fibril using Autodock 4.2.

**Proximity ligation assay.** Binding of cGAS and cytoplasmic mtDNA in human and mouse brain sections was determined by a PLA kit (Sigma-Aldrich) according to the manufacturer’s instructions. The primary antibodies used in PLA included anti-TFAM antibody (MA5-16148, Invitrogen), anti-cGAS antibody (human/mouse-specific, Invitrogen). PLA signals were visualized using a Zeiss LSM 980 Airyscan 2 NLO Two-Photon Confocal Microscope and indicated by PLA red dots per cell.

**Isolation of mitochondria and cytosolic fraction.** We isolated mitochondria from the hippocampus of Alzheimer’s disease brains or hippocampus of mouse brains as described previously(1). Mitochondria were isolated from brain homogenates by centrifugation at 1,500g and 12,000g, then resuspended in isolation buffer with digitonin and further centrifuged. Mitochondria were purified using a mitochondrial fractionation kit (Abcam) and protein concentration was measured using the Bio-Rad DC protein assay (Bio-Rad).

**Mitochondrial swelling assay for mPTP.** Calcium-triggered mitochondrial swelling assay was performed according to the method(2) with the mitochondria from the cortex of 8-month-old nonTg and Tg mice treated with CsA, BIP-V5, or NSC15384 were suspended in swelling assay buffer and exposed to 100 μM CaCl₂ to initiate swelling. Mitochondrial swelling was monitored at 540 nm using a spectrophotometer (Amersham Biosciences Ultrospect 3100) for 12 minutes with 30-second intervals.

**Enzyme-linked immunosorbent assay.** Mouse hippocampal tissue was extracted with 50 mM guanidine, and cerebral Aβ40/42 levels were measured by ELISA (Invitrogen). IFN-β and 2′3′-cGAMP in primary neuron, astrocyte, and microglial lysates after Aβ or gAβ treatment were quantified using IFN-β (abcam) and 2′3′-cGAMP (Cayman Chemical) ELISA kits. Additionally, 8-OHdG levels in mouse cortical lysates were determined by competitive ELISA (abcam). G3P, DHAP, and H2O2 in neuronal lysates after Aβ or gAβ treatment were measured using assay kits for G3P, DHAP (Sigma), and H2O2 (Invitrogen), per manufacturer protocols.

**Tetramethylrhodamine Methyl Ester Staining.** WT hippocampal neurons from day 1 newborn mice at DIV 12 were treated with 1μM Aβ or 1μM gAβ or vehicle for 24 h for Tetramethylrhodamine methyl ester (TMRM) analysis by flow cytometry (CytoFLEX), as previously described (3).

**Mitochondrial respiration complex activity.** Mitochondrial respiration complex activity was measured using homogenates of the hippocampus or neuronal cells as previously described(4). Hippocampal tissue or cells were lysed in isolation buffer. Protein extracts (50 μg) were used to measure complex I (NADH: ubiquinone oxidoreductase) and IV (cytochrome c oxidase) activities. Complex I activity was assessed by absorbance change at 340 nm over 6 minutes. Complex IV activity was measured by absorbance at 550 nm using ferrocytochrome c as substrate, recorded at 10-second intervals for 1 minute.

**Measurement of ATP levels.** ATP levels were measured using an ATP Bioluminescence Assay Kit (Sigma/Roche). Mouse cortical tissues or neuronal cells were homogenized in the provided lysis buffer, incubated on ice for 30 minutes, and centrifuged at 12,000×g for 10 minutes at 4°C. ATP levels in the supernatant were quantified with a Luminescence plate reader (Molecular Devices), using a 1.6 s delay and 10 s integration time.

**Mitochondrial Electron paramagnetic resonance (EPR) measurements.** Evaluation of intracellular ROS levels was accessed by EPR spectroscopy(5). Neurons or brain tissues were incubated with CMH (cyclic hydroxylamine 1-hydroxy-3-methoxycarbonyl-2, 2, 5, 5-tetramethyl pyrrolidine, 100 μM) for 30 min, then washed three times with cold PBS. The brain tissues were collected and homogenized with 100 μL of PBS for EPR measurement. The EPR spectra were collected, stored, and analyzed with a Bruker EleXsys 540× -band EPR spectrometer (Billerica, MA, USA) using the Bruker software Xepr (Billerica, MA, USA).

**NanoString nCounter gene expression.** NanoString Mouse Neuroinflammation Panels were used to analyze neuropathology, microglial function, astrocyte activation, innate immunity, DNA damage, and cellular stress, with panel details provided in ***SI Appendix*, Table S3**. RNA was extracted from either the primary cell (neuronal, astrocytes, and microglia) or the CA1 region of the mouse brain hippocampus (CA1 region was poured from both hippocampi) using RNeasy Micro kits (Qiagen). RNA purity was confirmed by BioAnalyzer, and 100 ng/sample were hybridized with probes for 16 h at 65°C. The results obtained from the nCounter MAX Analysis System (NanoString Technologies, Seattle WA) were imported into the nSolver Analysis Software (v4.0; NanoString Technologies) for QC verification, normalization, and data statistical analysis using Advanced Analysis software (v2.0.115; NanoString Technologies). All assays were performed according to the manufacturer’s protocols.

**In situ spatial resolve analysis (Stereo-sequencing)**

It features a spatially resolved transcriptomic technology with high resolution and large field of view. Standard DNB chips have spots with approximately 220 nm diameter and a center-to-center distance of 500 or 715 nm **(fig. S12-IA)**, providing up to 400 spots for tissue RNA capture per 100 mm^2^. Frozen tissue sections (10mm thickness) are loaded on to the chip surface, followed by fixation, permeabilization to capture the tissue polyA-tailed RNA, and finally reverse transcription plus amplification. Amplified-barcoded cDNA is collected, used as template for library preparation, and sequenced together with the CID. Computational analysis of the sequencing data allows high-resolution spatially resolved transcriptomics known as spatial enhanced resolution omics-sequencing (Stereo-seq). To analyze the brain data, we performed unsupervised SCC of binned (bin 50, 50 x 50 DNB bins, 25 µm diameter) Stereo-seq data to identify the changes in disease associated MID count of genes with brain regions. We provide the details in SI Appendix.

**Generation of Stereo-seq chips.** To generate the patterned array, we synthesized two oligo sequences: one containing 25 random deoxynucleotides DNB library oligo1 and the other a fixed sequence with 5’ phosphorylated DNB library oligo2. These two oligos were ligated with splint oligo1 at 37°C for 2 hours using T4 ligase (NEB; 1U/ml T4 DNA ligase and 13 T4 DNA ligation buffer). The products were purified using the AMPure XP Beads (Vazyme, N411-03) and then PCR amplified with the following steps: 95°C for 5 minutes, 12 cycles at 98°C for 20 seconds, 58°C for 20 seconds, 72°C for 20 seconds and a final incubation at 72°C for 5 minutes. The PCR products were purified using the AMPure XP Beads. DNB were then generated by rolling circle amplification and loaded onto the patterned chips according to the MGI DNBSEQ-Tx sequencer manual. Next, to determine the distinct DNB-CID sequences at each spatial location, single-end sequencing was performed using a CID sequencing primer in MGI DNBSEQ-Tx sequencer with SE25 sequencing strategy. After sequencing, the capture oligo including 22 nt poly-T and 10 nt UMI was hybridized with the DNB in 5 x SSC buffer at 37°C for 30 minutes, and then incubated with T4 ligase (NEB, 1 U/ml T4 DNA ligase, 1 x T4 DNA ligation buffer and 0.5% PEG2000) at 37°C for 1 hour. This produces capture probes containing a 25 nt CID barcode, a 10 nt UMI and a 22 nt poly-T ready for poly-A RNA capture.

**Calling of CID.** CID sequences together with their corresponding coordinates for all DNB were determined using a base calling method according to manufacturer’s instruction of MGI DNBSEQ-Tx sequencer. After sequencing, the capture chip was split into smaller size chips (5 mm x 10 mm, 10 mm x 10 mm, 10 mm x 20 mm) ready for use. At this stage, we filtered out all duplicated CID that correspond to non-adjacent spots.

**Tissue processing.** Mouse brain sections were adhered to the Stereo-seq chip surface and incubated at 37^o^C for 3-5 minutes. Then, the sections were fixed in methanol and incubated for 40 minutes at -20C before Stereo-seq library preparation. Where indicated, the same sections were stained with nucleic acid dye (Thermo fisher, Q10212) and imaging was performed with a Ti-7 Nikon Eclipse microscope prior to in situ capture at the channel of FITC.

**In situ reverse transcription.** After washed with 0.1 3 SSC buffer (Thermo, AM9770) supplemented with 0.05 U/ml RNase inhibitor (NEB, M0314L), tissue sections placed on the chip were permeabilized using 0.1% pepsin (Sigma, P7000) in 0.01 M HCl buffer, incubated at 37°C for 12 minutes and then washed with 0.1 3 SSC buffer (Thermo, AM9770) supplemented with 0.05 U/ml RNase inhibitor (NEB, M0314L). RNA released from the permeabilized tissue and captured by the DNB was reverse transcribed overnight at 42C using SuperScript II (Invitrogen, 18064-014, 10 U/ml reverse transcriptase, 1 mM dNTPs, 1 M betaine solution PCR reagent, 7.5 mM MgCl2, 5 mM DTT, 2 U/ml RNase inhibitor, 2.5 mM Stereo-seq-TSO and 1 x First-Strand buffer). After reverse transcription, tissue sections were washed twice with 0.1 3 SSC buffer and digested with Tissue Removal buffer (10 mM Tris-HCl, 25 mM EDTA, 100 mM NaCl, 0.5% SDS) at 37C for 30 minutes. cDNA-containing chips were then subjected to Exonuclease I (NEB, M0293L) treatment for 1 hour at 37°C and were finally washed once with 0.1x SSC buffer.

**Amplification.** The resulting cDNAs were amplified with KAPA HiFi Hotstart Ready Mix (Roche, KK2602) with 0.8 mM cDNA-PCR primer. PCR reactions were conducted as follows: incubation at 95°C for 5 minutes, 15 cycles at 98°C for 20 seconds, 58°C for 20 seconds, 72°C for 3 minutes and a final incubation at 72^o^C for 5 minutes.

**Library construction and sequencing.** The concentrations of the resulting PCR products were quantified by Qubit dsDNA Assay Kit (Thermo, Q32854). A total of 20 ng of DNA were then fragmented with in-house Tn5 transposase at 55°C for 10 minutes, after which the reactions were stopped by the addition of 0.02% SDS and gently mixing at 37°C for 5 minutes after fragmentation. Fragmented products were amplified as described below: 25 ml of fragmentation product, 1 3 KAPA HiFi Hotstart Ready Mix and 0.3 mM Stereo-seq-Library-F primer, 0.3 mM Stereo-seq-Library-R primer in a total volume of 100 ml with the addition of nuclease-free H2O. The reaction was then run as: 1 cycle of 95°C 5 minutes, 13 cycles of 98°C 20 seconds, 58°C 20 seconds and 72°C 30 seconds, and 1 cycle of 72°C 5 minutes. PCR products were purified using the AMPure XP Beads (0.63 and 0.153), used for DNB generation and finally sequenced on MGI DNBSEQ-Tx sequencer with read length.

**Stereo-seq raw data processing.** Fastq files were generated using a MGI DNBSEQ-Tx sequencer. CID and MID are contained in the read 1 (CID: 1-25 bp, MID: 26-35 bp) while the read 2 consist of the cDNA sequences. CID sequences on the first reads were first mapped to the designed coordinates of the in situ captured chip achieved from the first round of sequencing, allowing 1 base mismatch to correct for sequencing and PCR errors. Reads with MID containing either N bases or more than 2 bases with quality score lower than 10 were filtered out. CID and MID associated with each read were appended to each read header. Retained reads were then aligned to the reference genome (mm10) using STAR (Dobin et al., 2013) and mapped reads with MAPQ > 10 were counted and annotated to their corresponding genes). UMI with the same CID and the same gene locus collapsed, allowing 1 mismatch to correct for sequencing and PCR errors. Finally, this information was used to generate a CID-containing expression profile matrix. The whole procedure was integrated into a publicly available pipeline SAW available at <https://github.com/BGIResearch/SAW>.

**Mice behavior studies.** Mice behavior was assessed using the Morris Water Maze (MWM) and Nest behavior test after four weeks of inhibitor administration. Spatial learning and memory were investigated with the Morris Water Maze (MWM) and Nest behavior test were performed after four weeks of inhibitor administration. Mice were subjected to the MWM hidden platform test as described previously(6). Mice were trained for five days with four trials daily and assessed for spatial memory on the final day using a probe trial. Traces were recorded and analyzed with HVS Image 2014, with investigators blinded to mouse identities. For nesting behavior, mice were given a pre-weighed Nestlet (3.0 g; Ancare) for 24 hours, and nesting capability was assessed by measuring the remaining cotton. Nest quality was scored on a scale of 1 to 5, according to our previously published criteria(5), with 1 being largely untorn nestlets and 5 being near perfect nests.

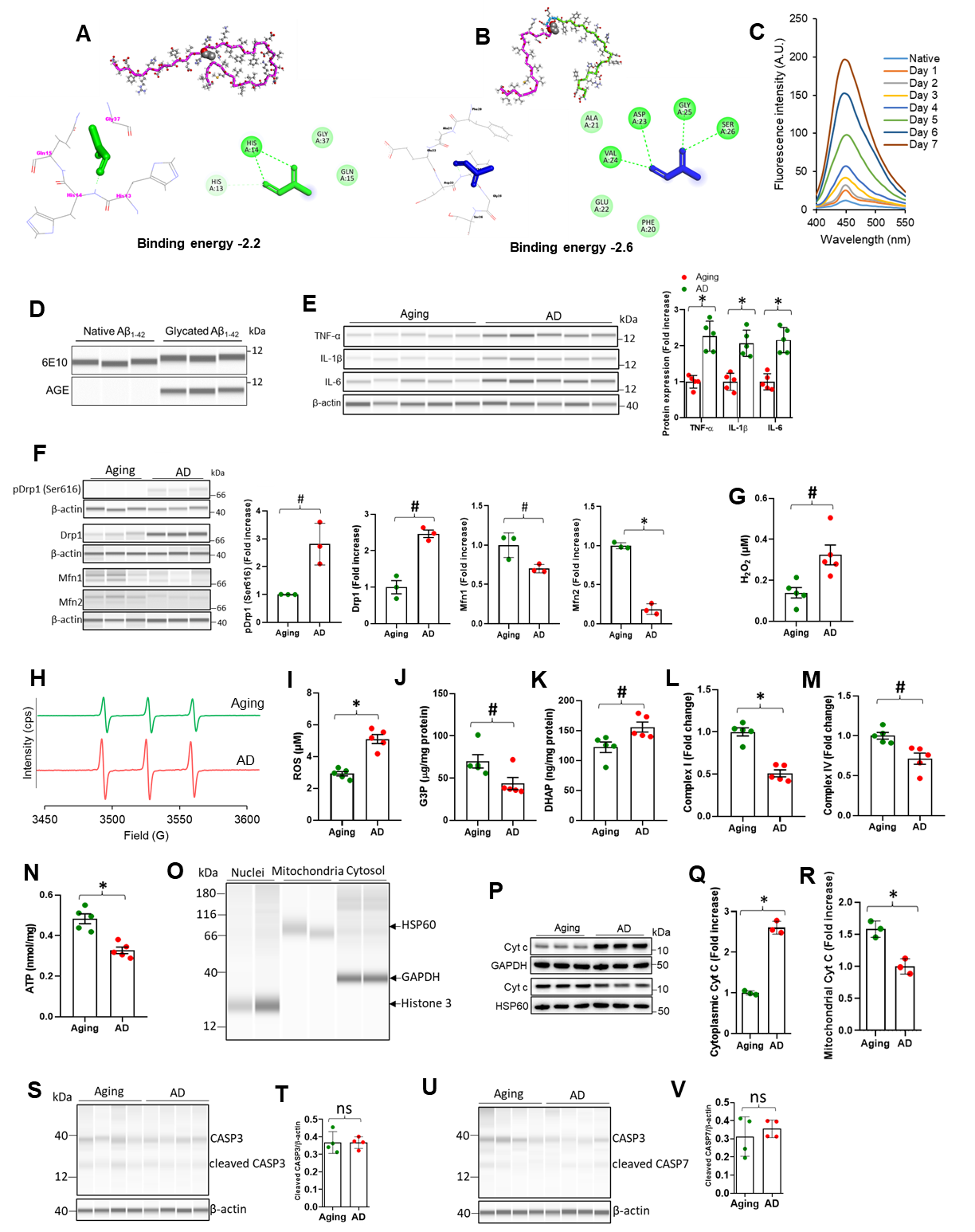

**Fig. S.1. Interaction of methylglyoxal with Aβ, mitochondrial dysfunction and cGAS-STING activation in human AD brain. (A, B)**, The structure of Aβ (40 and 42 residues) was obtained from the protein data bank using the PDB IDs 2M4J (40 residues) and 8eze42 (42 residues). The PubChem database was accessed to find the 3D conformer of methylglyoxal (MGO). These structures were prepared for docking analysis using Discovery Studio 2021, and the prepared methylglyoxal was docked with both the structure Aβ40 (a) and Aβ42 (b) residues of Aβ fibril using Autodock 4.2. **(C)**, fluorescence emission spectra of Aβ42 (native, Day 8) and Aβ42-MGO (Day 1- Day 7). The excitation wavelength was 350 nm, and the emission wavelength was 450 nm. **(D)** In vitro–glycated synthetic Aβ_1-42_ was detected by immunoblotting using anti-6E10 antibody, followed by re-probing with anti-AGE antibodies; n = 3/group. (**E**), Immunoblotting analysis of the expression and quantification of TNF-α, IL-1β and IL-6, relative to β-actin in hippocampus region of aging or AD samples. Mean ± SD; n = 5; *P < 0.001, two-tailed Student’s *t*-test. (**F**), Immunoblotting analysis of the expression and quantification of pDrp1 (Ser616), Drp1, Mfn1, and Mfn2 relative to β-actin in aging and AD human brain cortical samples. Mean ± SD; n = 3; ^#^P < 0.05, *P < 0.01, one-way ANOVA with Bonferroni’s post hoc test. (**G**), Levels of H_2_O_2_ was determined in indicated aging or AD human brain cortical lysates. Mean ± SD; n = 5; ^#^P < 0.05. one-way ANOVA with Bonferroni’s post hoc test. (**H, I**), Representative and quantified *in vitro* EPR spectra measured in aging and AD human cortical tissue lysates are shown. The peak height in the spectrum indicates levels of ROS. Mean ± SD; n = 5; *P < 0.01. one-way ANOVA with Bonferroni’s post hoc test. (**J-N**), Levels of, G3P (J), DHAP (K), activity of complex I (L), complex IV (M), and ATP level (N) were determined in indicated aging or AD human brain cortical lysates. Mean ± SD; n = 5; ^#^P < 0.05, *P < 0.01. one-way ANOVA with Bonferroni’s post hoc test. (**O**) Representative immunoblot showing the purity of nuclear, mitochondrial, and cytosolic fractions. (**P-R**), Immunoblotting analysis of the expression and quantification of cytochrome c (Cyt c) relative to GAPDH or HSP60 in cytosolic and mitochondrial fraction respectively of the cortical aging or AD samples. Mean ± SD; n = 3 ;*P < 0.01. One-way ANOVA with Bonferroni’s post hoc test. (**S, T**), Immunoblotting analysis of the expression and quantification of Cleaved CASP3 (Caspase 3) relative to β-actin of the cortical aging or AD samples. Mean ± SD; n = 4 ^ns^P<0.05. One-way ANOVA with Bonferroni’s post hoc test. (**U, V**), Immunoblotting analysis of the expression and quantification of Cleaved CASP7 (Caspase 7) relative to β-actin of the cortical aging or AD samples. Mean ± SD; n = 4 ^ns^P<0.05. One-way ANOVA with Bonferroni’s post hoc test.

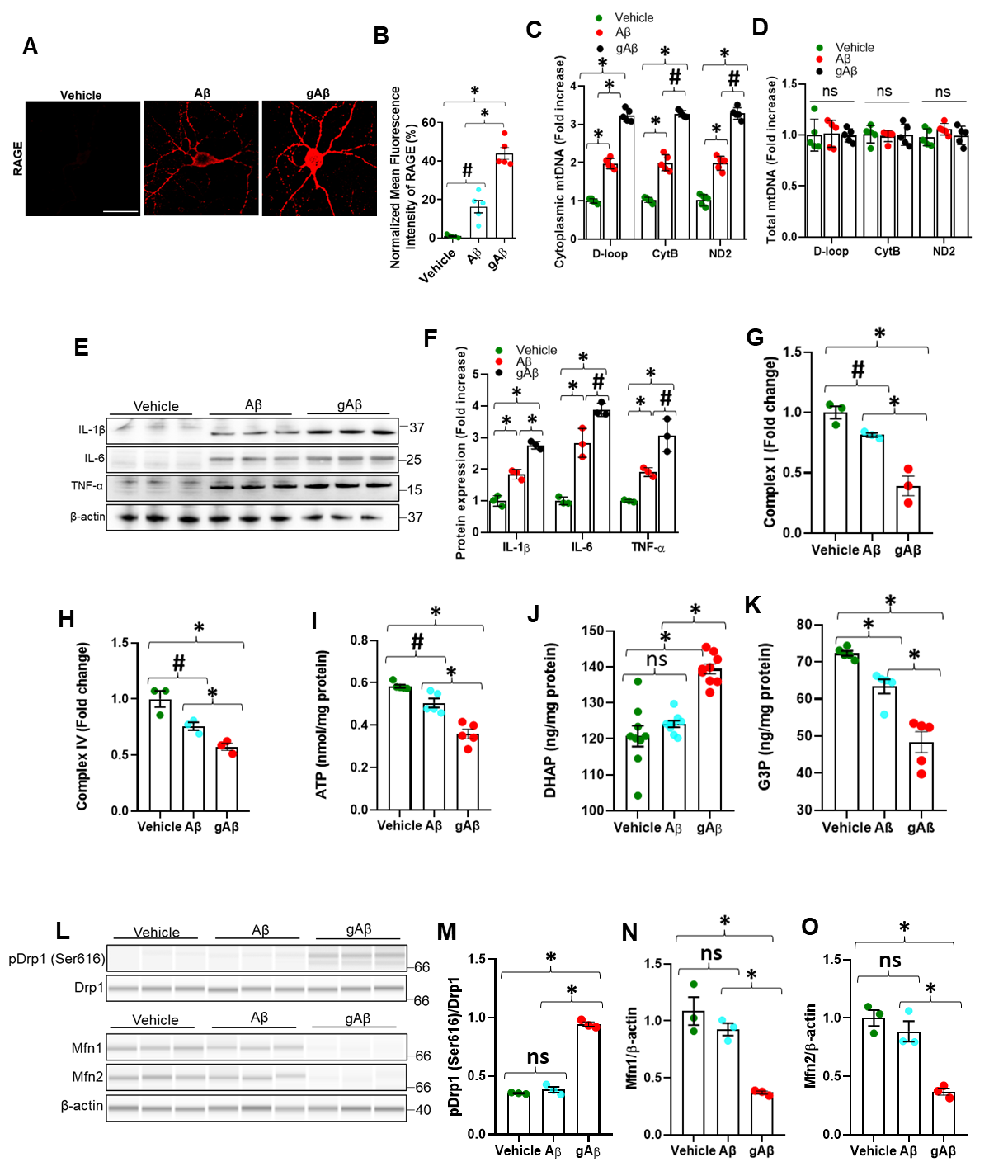

**Fig. S.2. Neuroinflammation, mitochondrial dysfunction, and change in mitochondrial dynamics after Aβ or gAβ treatment. (A),** Immunostaining of RAGE in hippocampal neurons (DIV 10) treated with vehicle, Aβ or gAβ for 24 h. Scale bar, 50 μm. Representative images as shown were from the results of five samples in each group. (**B**) Percentage AGE colocalization with Amyloid plaque. Mean ± SD; n = 5; ^#^P < 0.05, *P < 0.01, one-way ANOVA with Bonferroni’s post hoc test. (**C, D**), Quantification of cytoplasmic and total mitochondrial DNA (mtDNA) by qPCR in indicated cultured neurons with vehicle, Aβ or gAβ treatment for 24 h. D-loop, CytB and ND2 are specific fragments or proteins encoded by the mitochondrial genome. Mean ± SD; n = 5; *P < 0.01, ^#^P < 0.05, or ^ns^P > 0.05, one-way ANOVA with Bonferroni’s post hoc test. (**E, F**), Immunoblotting analysis of the expression and quantification of TNF-α, IL-1β and IL-6, relative to β-actin in indicated neurons (DIV 10) with the same treatment conditions. Mean ± SD; n = 3; ^#^P < 0.05, *P < 0.001, one-way ANOVA with Bonferroni’s post hoc test. (**G-I**), Activity of complex I (G), complex IV (H), and ATP level (I) were determined in indicated neuronal lysates with the same treatment condition. Mean ± SD; n = 3-5; ^ns^P > 0.05, ^#^P < 0.05, *P < 0.01. one-way ANOVA with Bonferroni’s post hoc test. (**J, K**), Activity of DHAP (J) and G3P (K) were determined in primary neuron (DIV 10) treated with vehicle, Aβ or gAβ. Mean ± SD; n = 3-9; ^ns^P > 0.05, *P < 0.01; ns: not significant. one-way ANOVA with Bonferroni’s post hoc test. (**L-O**), Immunoblotting analysis of the expression and quantification of pDrp1 relative to Drp1 and Mfn1, Mfn2 relative to β-actin in indicated neurons (DIV 10) with the same treatment conditions. Mean ± SD; n = 3; ^ns^P > 0.05, *P < 0.001, one-way ANOVA with Bonferroni’s post hoc test.

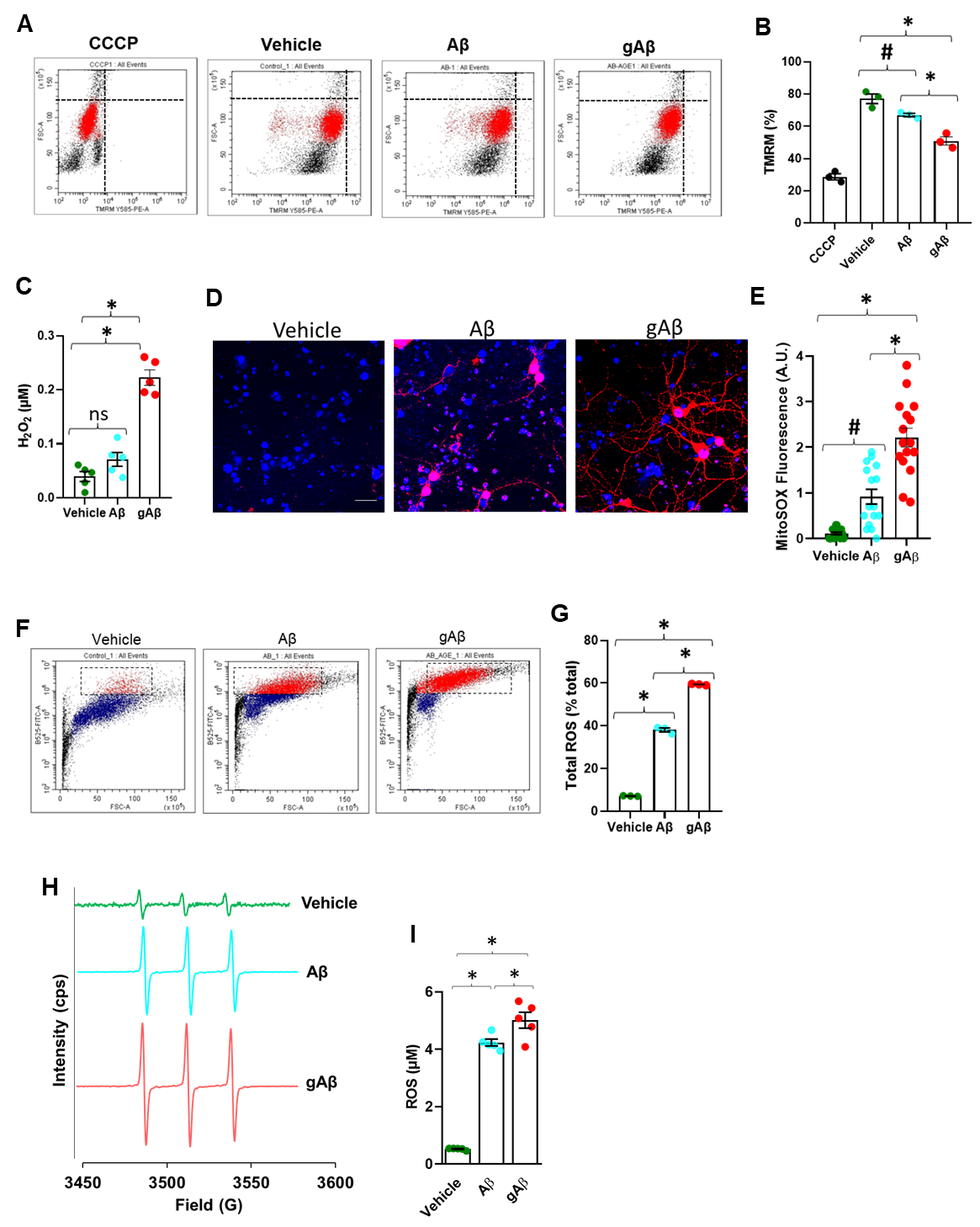

**Fig. S.3. Mitochondrial stress and neuropathology after Aβ or gAβ treatment.** (**A, B**), Mitochondrial membrane potential (TMRM) was estimated in indicated neurons (DIV 11) with the same treatment condition by FACS analysis. CCCP was used as positive control. Mean ± SD; n = 3; ^#^P < 0.05, **P* < 0.01. one-way ANOVA with Bonferroni’s post hoc test. (**C**), Level of H_2_O_2_ were determined in primary neuron (DIV 10) treated with vehicle, Aβ or gAβ. Mean ± SD; n = 5; ^ns^P > 0.05, *P < 0.01; ns: not significant. one-way ANOVA with Bonferroni’s post hoc test. (**D**) Mitochondrial superoxide levels were analyzed using the fluorescent dye MitoSox Red in indicated neurons (DIV 12) with the same treatment conditions. (**E**) Quantification of fluorescence intensity and data are mean ± SD. n = 3 images/field, ^#^P <0.01, *P <0.01. one-way ANOVA with Bonferroni’s post hoc test. (**F, G**) Total cellular ROS were estimated in indicated neurons (DIV 11) with the same treatment condition by FACS analysis. Mean ± SD; n = 3; *P < 0.01. one-way ANOVA with Bonferroni’s post hoc test. (**H, I**), Representative *in vitro* EPR spectra measured in primary neuron (DIV 10) treated with vehicle, Aβ or gAβ are shown. The peak height in the spectrum indicates levels of ROS (H). Quantification of EPR spectra in the indicated neuronal lysate (I). *P <0.01; Mean ± SD; n = 5; one-way ANOVA with Bonferroni’s post hoc test.

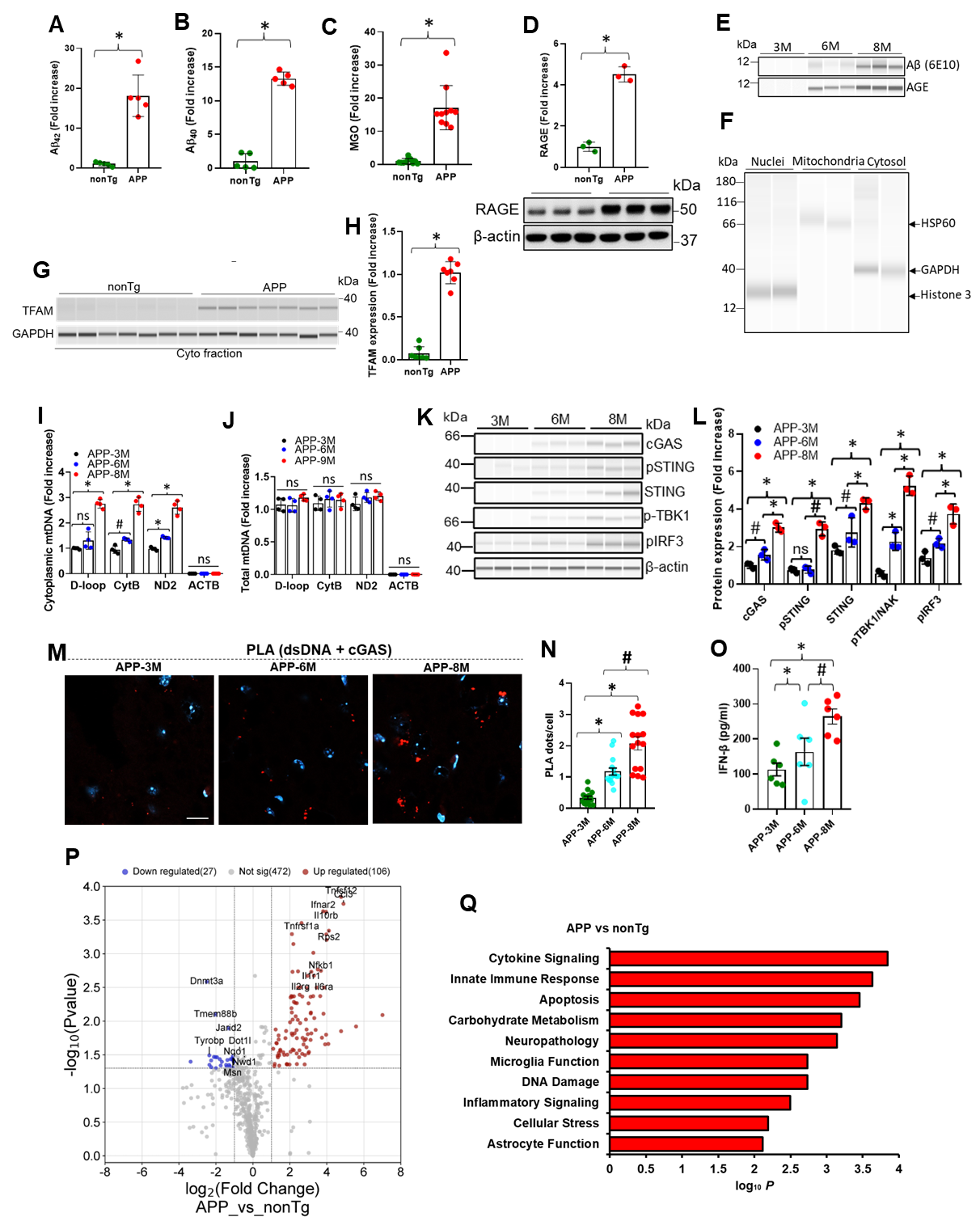

**Fig. S.4. Glycated Aβ stress and cGAS-IFN response in APP mice.** (**A, B**), Quantification of amyloid beta 42 (Aβ42) (a) and Aβ40 (b) in brain cells of 8-month-old nonTg and APP mice. Mean ± SD; n = 5; *P<0.001, two-tailed Student’s t-test. (**C**), Quantification of methylglyoxal (MGO) by ELISA in hippocampal cells of 8-month-old APP mice. Mean ± SD; n = 10; *P < 0.001, two-tailed Student’s *t*-test. (**D**), Immunoblotting analysis of the expression of RAGE and quantification in the indicated mice. Mean ± SD; n = 3; *P < 0.001, two-tailed Student’s *t*-test. (**E**) The cortical extracts from the brains of 3, 6, and 8-month-old APP mice were immunoprecipitated with Aβ antibody (6E10) and AGE. Representative immunoblots show the presence of the Aβ-AGE (gAβ) complex in the indicated mouse brain cells. (**F**) Representative immunoblot showing the purity of nuclear, mitochondrial, and cytosolic fractions. (**G, H**), Immunoblotting analysis of the expression (G), and quantification (H) of TFAM in cytosolic fraction of the indicated mice relative to GAPDH. Mean ± SD; n = 7; *P < 0.001, two-tailed Student’s *t*-test. (**I, J**), Quantification of cytoplasmic and total mtDNA by qPCR in brain cells of 3, 6, and 8-month-old APP mice. D-loop, CytB, and ND2 are specific fragments or proteins encoded by the mitochondrial genome. Mean ± SD; n = 5; ^ns^P>0.05, ^#^P <0.05, *P <0.01, ns: not significant, one-way ANOVA with Bonferroni’s post hoc test. (**K, L**), Expression and quantification of the expression of the cGAS, p-STING, p-TBK1, and p-IRF3 relative to β-actin. Mean ± SD; n = 3; ^ns^P > 0.05, ^#^P < 0.05, *P < 0.01, one-way ANOVA with Bonferroni’s post hoc test. (**M**), PLA with anti-dsDNA and anti-cGAS antibodies showing the association of cGAS protein with cytosolic or nuclear dsDNA in hippocampal tissues of 3, 6, and 8-month-old APP mice. Scale bar, 10 μm. (**N**), quantification of PLA red dots per cell. Mean ± SD; n = 5; five sections of each brain sample were stained and counted; *P < 0.001, ^#^P < 0.05, one-way ANOVA with Bonferroni’s post hoc test. (**O**), ELISA analysis of IFN-β level in the hippocampus of 3, 6, and 8-month-old APP mice. Mean ± SD; n = 6; *P < 0.001, or ^#^P < 0.05, one-way analysis of variance (ANOVA) with Bonferroni’s post hoc test. (**P**), The volcano plot shows significant DEGs (FDR ≤ 0.05, log_2_[FC] ≥ 0.3) in the hippocampus from APP versus nonTg (n=3). (**Q**) Top 10 Gene Ontology biological processes identified through analysis of the differentially expressed genes between APP and nonTg mice. DEGs, differentially expressed genes; FDR, false discovery rate; FC, fold change.

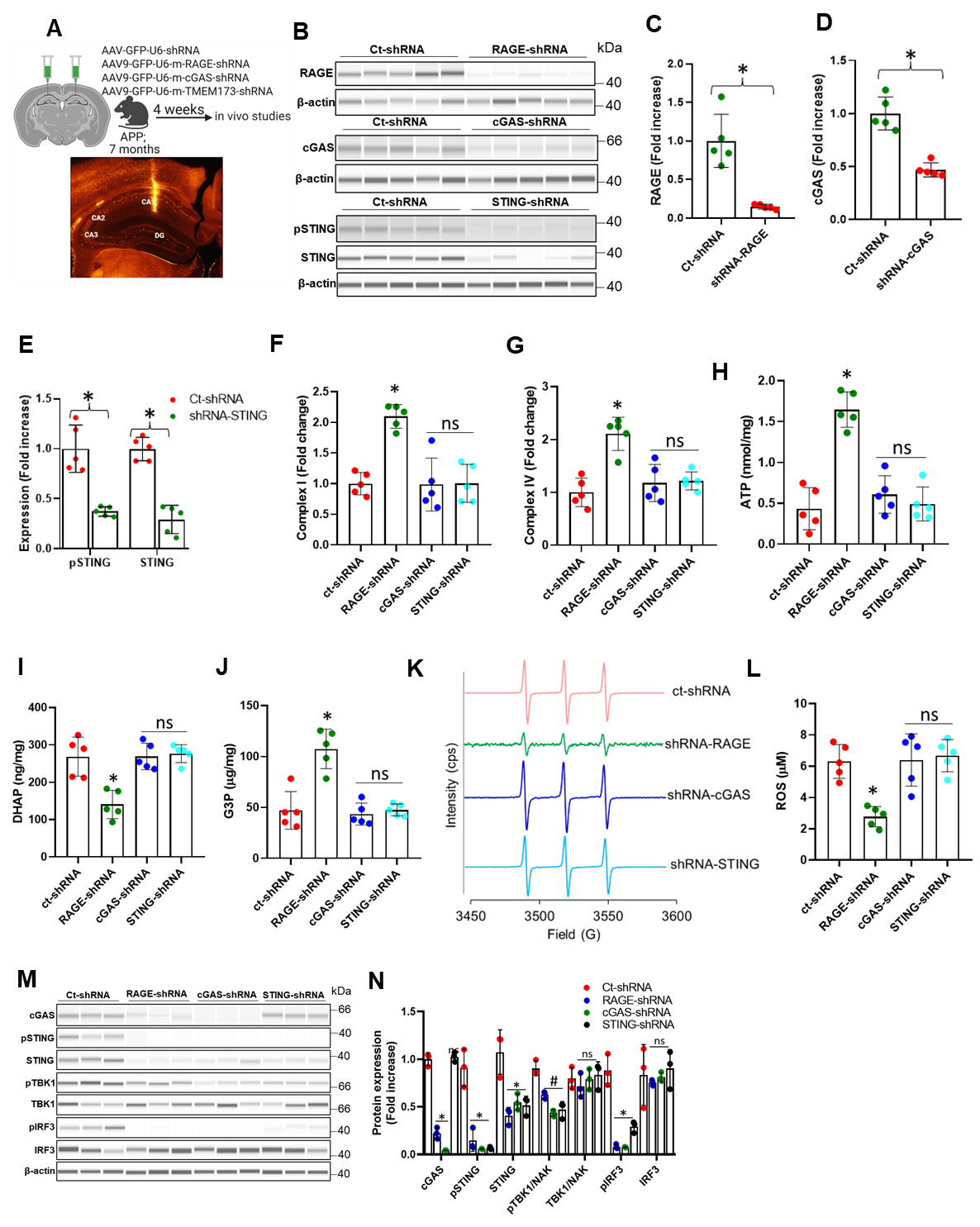

**Fig. S.5. Knockdown of RAGE, cGAS or STING restores mitochondrial functionality and attenuation oxidative stress.** (**A**) Schematic representation of stereotactic injection into the CA1 region of 7 months old APP mice’s hippocampus. (**B-E**), Immunoblotting analysis and quantification of the expression of the RAGE (C), cGAS (D), pSTING and STING (E) in hippocampal CA1 regions’ protein lysate of the indicated mice; n = 5; *P < 0.001, two-tailed Student’s *t*-test. (**F-J**), Activity of complex I (F), complex IV (G), and level of ATP (H), DHAP (I), and G3P (J) were determined in Ct-shRNA, RAGE-shRNA, cGAS-shRNA, and STING-shRNA transfected 8-month-old APP mice hippocampal CA1 region lysate (both hippocampal CA1 regions were poured together). Mean ± SD; n = 5; ^ns^P > 0.05, *P < 0.001; ns: not significant. one-way ANOVA with Bonferroni’s post hoc test. (**K, L**), Representative peak height in the in vitro EPR spectrum indicating levels of ROS and quantification of the EPR spectra measured in indicated samples of Ct-shRNA, RAGE-shRNA, cGAS-shRNA, and STING-shRNA transfected APP mice lysates. Mean ± SD; n = 5; ^ns^P > 0.05, *P < 0.01. one-way ANOVA with Bonferroni’s post hoc test. (**M**) Immunoblotting analysis of the expression of the cGAS, p-STING, STING, p-TBK1, TBK1, and p-IRF3, IRF3 in the indicated samples; n = 3. (**N**), Quantification of the expression of the cGAS, p-STING, STING, p-TBK1, TBK1, and p-IRF3, IRF3 relative to β-actin. Mean ± SD; n = 3; nsP > 0.05, #P < 0.05, *P < 0.001, one-way analysis of variance (ANOVA) with Bonferroni’s post hoc test.

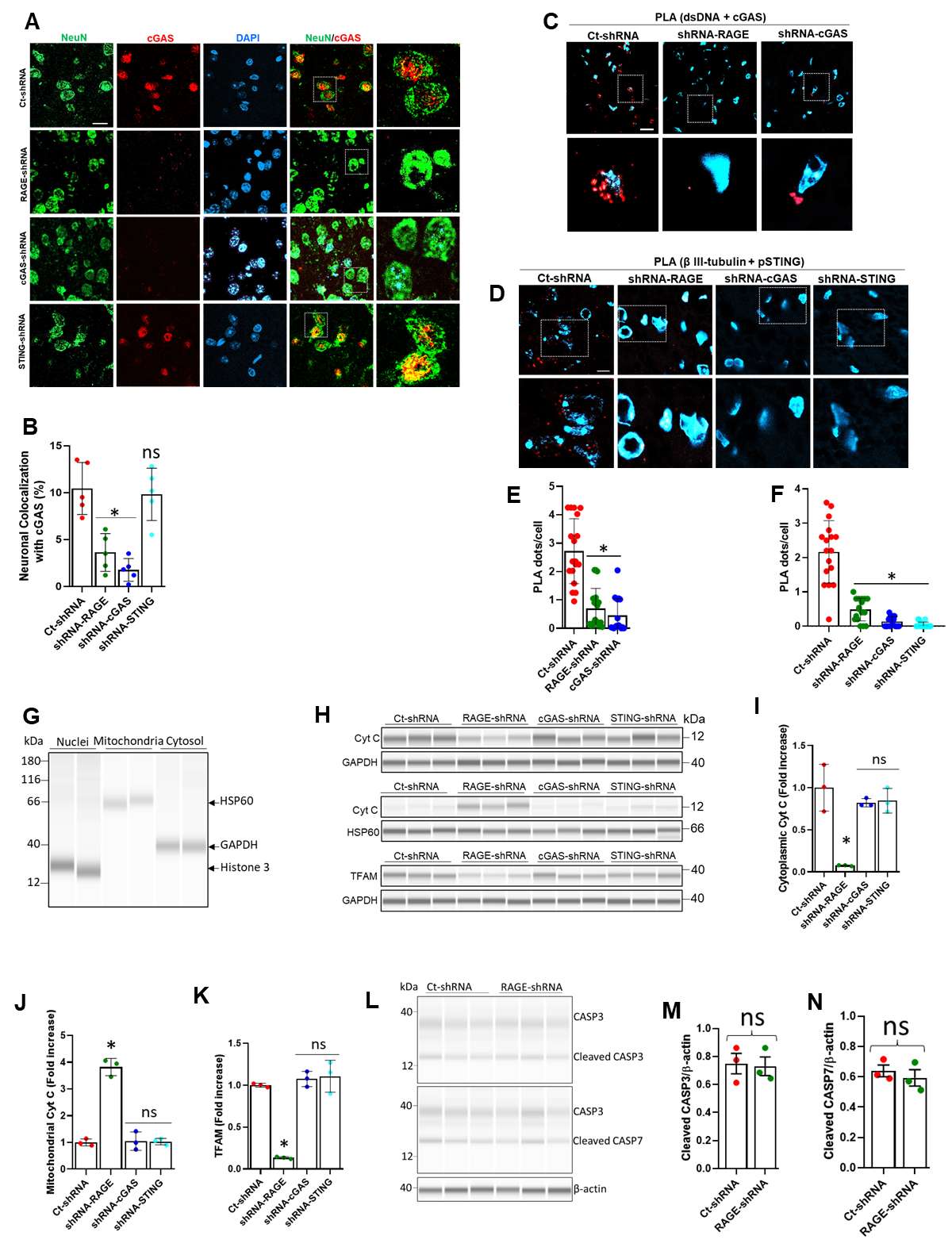

**Fig. S.6. Knockdown of RAGE, cGAS, or STING attenuates mtDNA and cytochrome c release.** (**A**), Immunostaining of cGAS with NeuN in the hippocampal CA1 region of indicated mice. Scale bar, 10 μm. (**B**) Percentage neuronal colocalization with cGAS. Mean ± SD; n = 5; ^ns^P > 0.05, *P < 0.001, one-way ANOVA with Bonferroni’s post hoc test. (**C, D**), PLA with anti-dsDNA and anti-cGAS antibodies or anti-β III-tubulin and anti-pSTING antibodies showing the association of cGAS protein with cytosolic or nuclear dsDNA or neuronal protein with pSTING in hippocampal CA1 region of the indicated mice. Scale bar, 10 μm. (**E, F**) Quantification of PLA red dots per cell. Mean ± SD; n = 5; ^ns^P > 0.05, *P < 0.001, one-way ANOVA with Bonferroni’s post hoc test. (**G**) Representative immunoblot showing the purity of nuclear, mitochondrial, and cytosolic fractions. (**H**), Immunoblotting analysis of the expression of the cytochrome c (Cyt c) in the indicated samples cytosolic and mitochondrial fraction and TFAM in the cytosol. (**I, J**), Quantification of the expression of the CytC relative to GAPDH or HSP60 in cytosolic and mitochondrial fraction respectively. Mean ± SD; n = 3; ^ns^P > 0.05, *P < 0.001, one-way ANOVA with Bonferroni’s post hoc test. (**K**), Quantification of the TFAM relative to GAPDH in cytosolic fraction. Mean ± SD; n = 3; ^ns^P > 0.05, *P < 0.001, one-way ANOVA with Bonferroni’s post hoc test. (**L-N**), Immunoblotting analysis of the expression and quantification of Cleaved CASP3 and CASP7 relative to β-actin in the indicated samples of Ct-shRNA and RAGE-shRNA transfected APP mice lysates. Mean ± SD; n = 3 ^ns^P<0.05. One-way ANOVA with Bonferroni’s post hoc test.

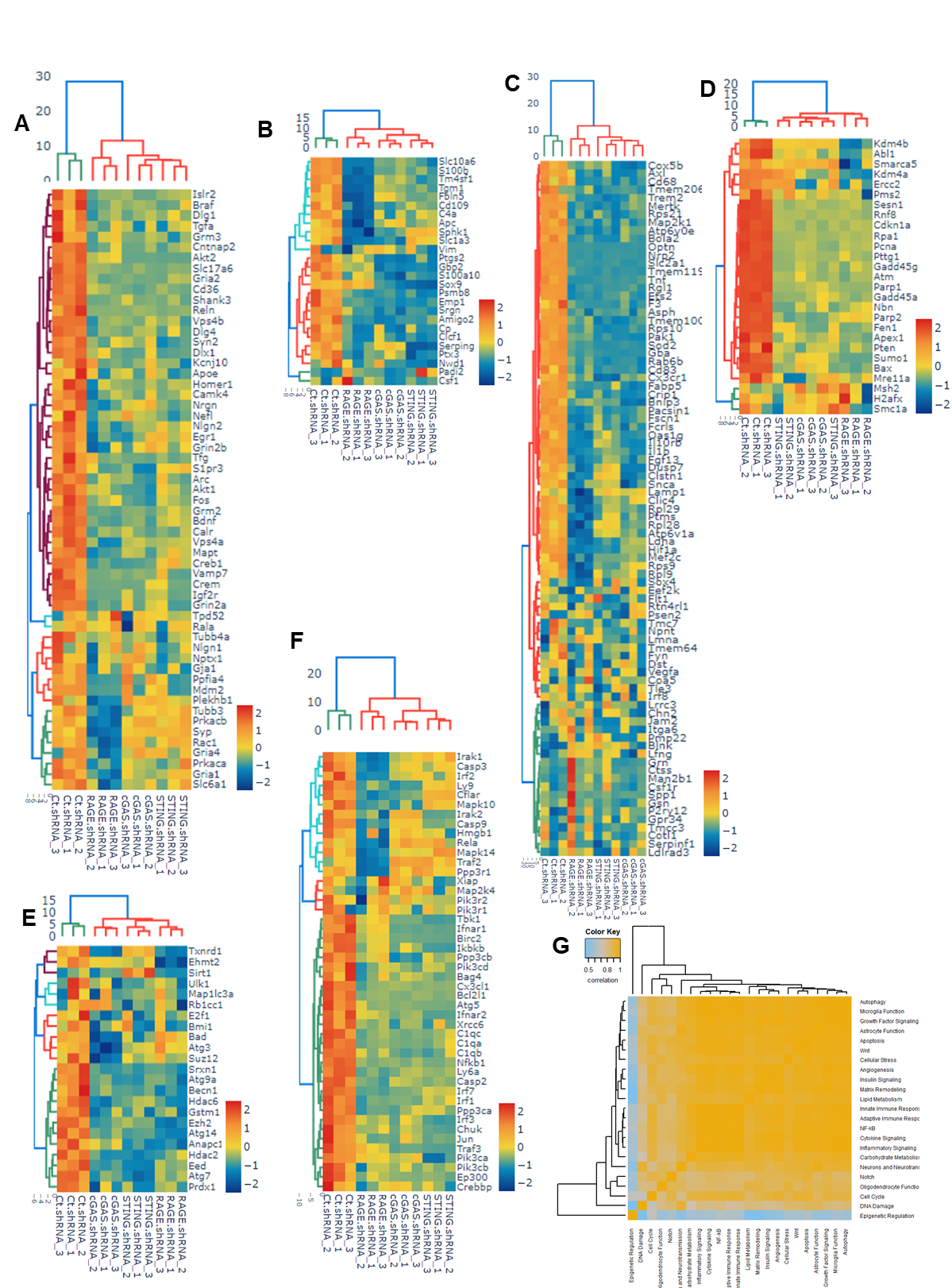
**Fig. S.7. Knockdown of RAGE, cGAS, or STING downregulates neuropathological genes and attenuates innate immune response.** (**A-F**), NanoString nCounter neuroinflammation profiling panels were employed to analyze extracted RNA from knockdown of RAGE, cGAS, STING, or control mice’s CA1 hippocampi (pouring both hippocampi regions for RNA extraction) regions. The gene expression heatmap shows NanoString analysis of neuropathology (A), astrocyte function (B), microglial function (C), innate immune response (D), DNA damage (E), and cellular stress (F) annotation of the indicated extracted RNA (n = 3). (**G**), Representation of heatmap of correlation matrix of pathway scores of all samples from NanoString nCounter analysis.

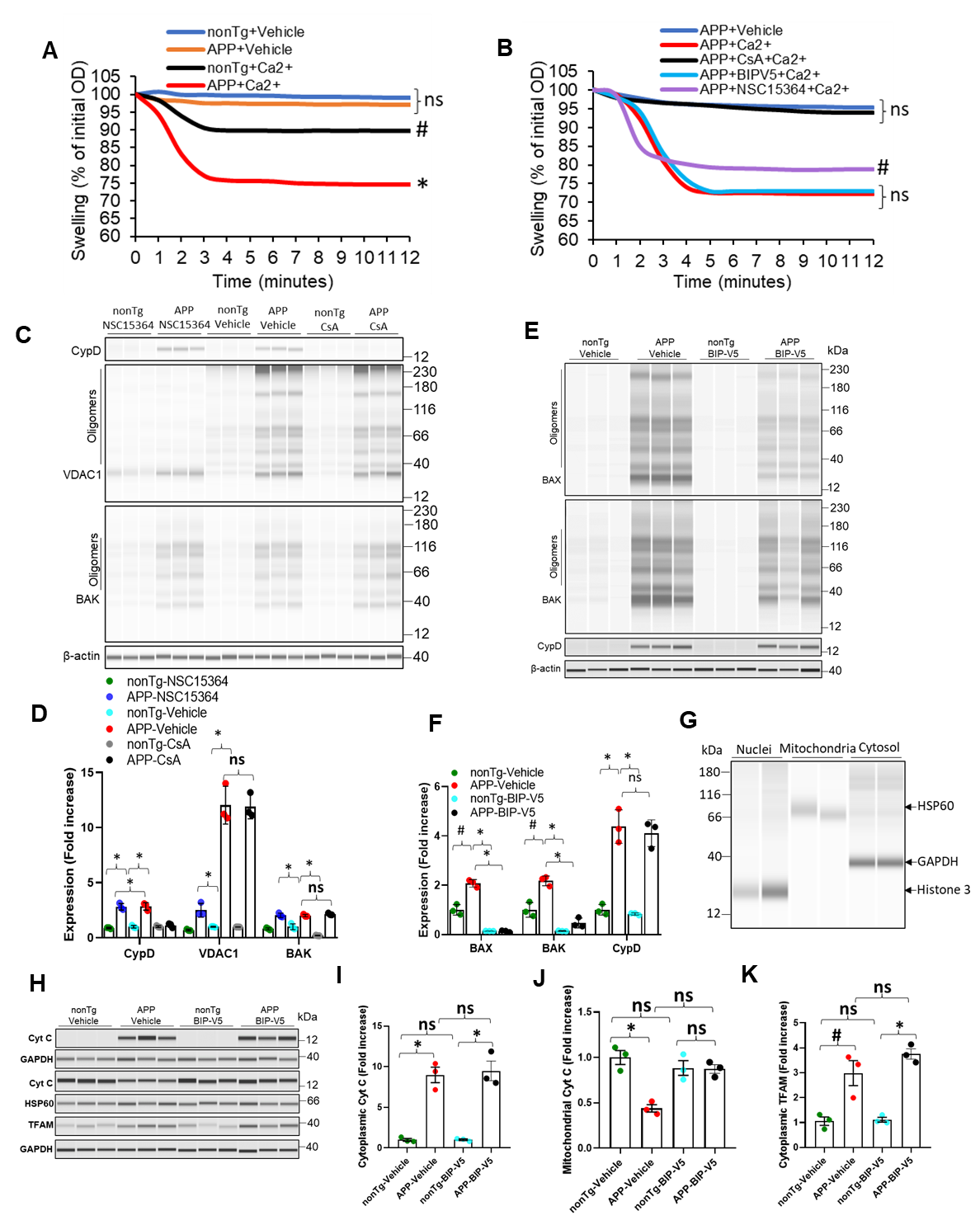

**Fig. S.8. Relation of VDAC1 to CypD and BAX, and inhibition of VDAC1 restrains mtDNA and cytochrome c leakage.** (**A**) Representative results of swelling from the nonTg or APP mice cortical mitochondria (8 months old). Data are shown as the percentage change relative to the initial OD at an absorbance of 540 nm. ^#^P < 0.05 versus Ca^2+^ (100μM)-induced nonTg cortical mitochondria or *P < 0.01 versus Ca^2+^ (100μM)-induced APP cortical mitochondria. Mean ± SD; n = 5; one-way ANOVA with Bonferroni’s post hoc test. (**B**), Representative results of calcium uptake in cortical mitochondria from the vehicle, CsA, BIP-V5 or NSC15364 administered 8 months old APP mice or vehicle treated 8-months old nonTg mice. Mitochondrial swelling was induced by Ca^2+^ (100 μM) and the data are shown as the percentage change relative to the initial OD at an absorbance of 540 nm. ^ns^P > 0.05, *^#^*P *< 0.05 or **P *<* 0.01 versus Ca^2+^ (100μM)-induced APP cortical mitochondria. Mean ± SD; n = 5; one-way ANOVA with Bonferroni’s post hoc test. (**C**) Immunoblotting analysis of the expression of the CypD, VDAC1 and BAK in hippocampus of NSC15364 or CsA administered APP mice; n = 3. (**D**), Quantification of the expressions of the CypD, BAX and VDAC1 relative to β-actin. Mean ± SD; n = 3; ^ns^P > 0.05, *P < 0.001, one-way analysis of variance (ANOVA) with Bonferroni’s post hoc test. (**E**) Immunoblotting analysis of the expression of the BAX, BAK and CypD in hippocampus of BIP-V5 administered APP mice; n = 3. (**F**), Quantification of the expression of the BAX, BAK and CypD relative to β-actin. Mean ± SD; n = 3; ^ns^P > 0.05, ^#^P < 0.05, *P < 0.001, one-way ANOVA with Bonferroni’s post hoc test. (**G**) Representative immunoblot showing the purity of nuclear, mitochondrial, and cytosolic fractions. (**H**), Immunoblotting analysis of the expression of the Cyt c in the BIP-V5 administered APP hippocampal cytosolic and mitochondrial fraction; n = 3. Immunoblotting analysis for the TFAM in the indicated samples cytosolic fraction. n = 3. (**I, J**), Quantification of the expression of the CytC relative to GAPDH or HSP60 in cytosolic and mitochondrial fraction respectively. Mean ± SD; n = 3; ^ns^P > 0.05, **P* < 0.001, one-way ANOVA with Bonferroni’s post hoc test. (**K**), Quantification of the TFAM relative to GAPDH in cytosolic fraction. Mean ± SD; n = 3; ^ns^P > 0.05, **P* < 0.001, one-way analysis of variance (ANOVA) with Bonferroni’s post hoc test.

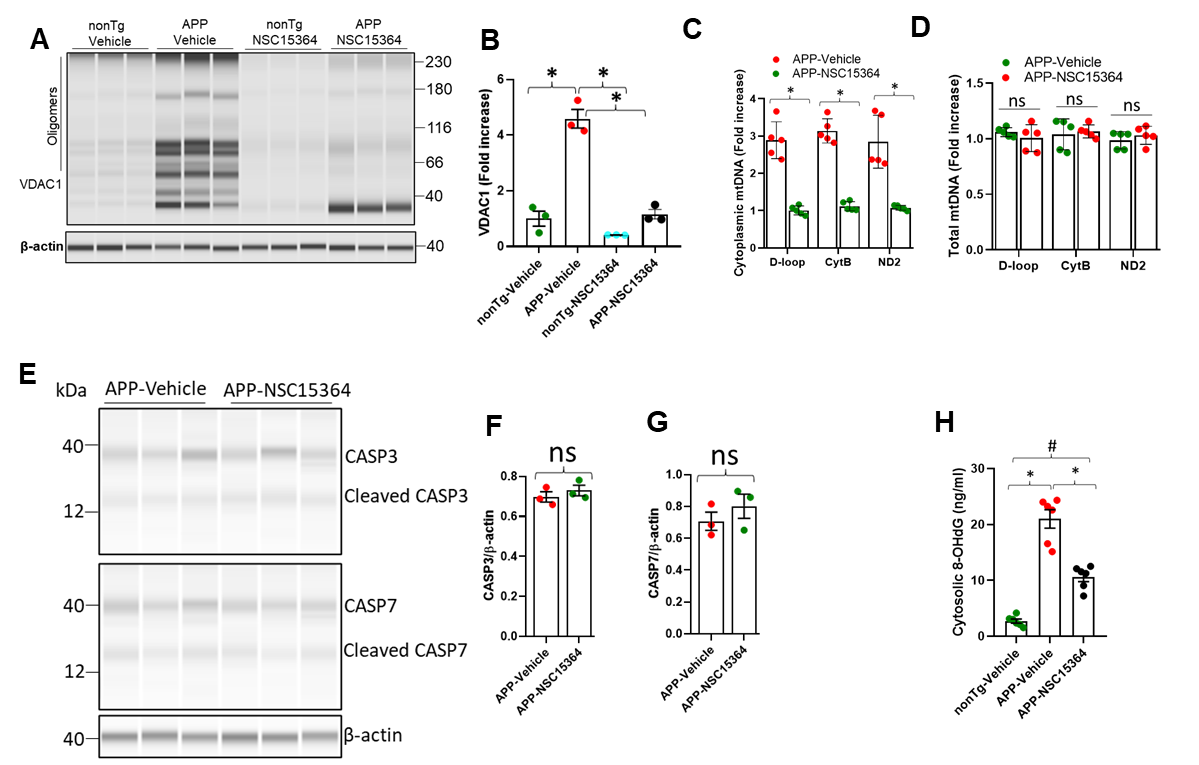

**Fig. S.9. VDAC1 inhibition alleviates mtDNA leakage and related stress in APP mice.** (**A**), Immunoblotting analysis of the expression of the VDAC1 in hippocampus of NSC15364 administered APP mice; n = 3. (**B**), Quantification of the expression of the VDAC1 relative to β-actin. Mean ± SD; n = 3; *P < 0.001, ANOVA with Bonferroni’s post hoc test. (**C**), Quantification of cytoplasmic mtDNA by qPCR in brain cortical cells of NSC15364 treated 8-month-old APP mice. (**D**), Quantification of total mtDNA by qPCR in brain cortical cells of NSC15364 treated 8-month-old APP mice. D-loop, CytB, and ND2 are specific fragments or proteins encoded by the mitochondrial genome. Mean ± SD; n = 5; ^ns^P>0.05, *P<0.001, ns, not significant, two-tailed Student’s *t*-test. (**E-G**), Immunoblotting analysis of the expression and quantification of Cleaved CASP3 and CASP7 relative to β-actin in hippocampus of NSC15364 administered APP mice; n = 3. ^ns^P<0.05. two-tailed Student’s *t*-test. (**H**), Quantification of 8-OHdG (ng/ml) by enzyme-linked immunosorbent assay (ELISA) in cortical brain (including hippocampus) cytosolic fraction of nonTg (vehicle-treated) and APP (NSC15364-treated) mice. Mean ± SD; n = 6; ^#^P < 0.05, *P < 0.001, one-way ANOVA with Bonferroni’s post hoc test.

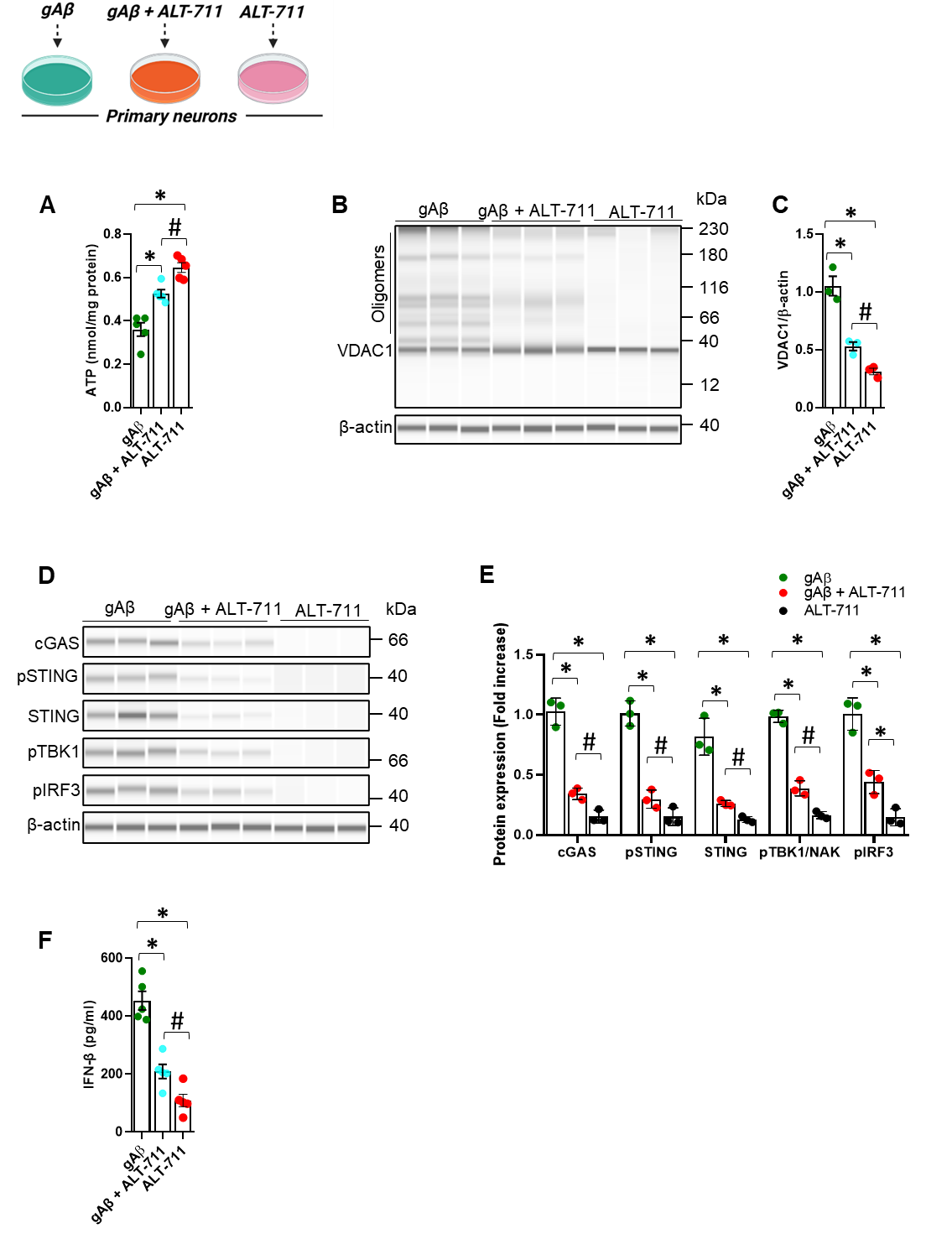

**Fig. S.10. ALT-711 mitigates** **glycated Aβ-induced mitochondrial dysfunction and cGAS-STING activation in neurons.** (**A**), ATP levels were measured in cortical primary neurons (DIV 10) treated with 5μM glycated Aβ (gAβ) with/without ALT-711 (20μM) for 24hourss. Mean ± SD; n = 5; P < 0.05, *P < 0.01. one-way ANOVA with Bonferroni’s post hoc test. **(B, C)**, Immunoblotting analysis of the expression and quantification of VDAC1 relative to β-actin in indicated neurons (DIV 10) with the same treatment conditions. Mean ± SD; n = 3; #P < 0.05, *P < 0.01, one-way ANOVA with Bonferroni’s post hoc test. (**D, E**), Immunoblot analysis for the expression and quantification of the expression of the cGAS, p-STING, STING, p-TBK1, and p-IRF3, relative to β-actin in the indicated neuronal lysate. Mean ± SD; n = 3; ^#^P < 0.05, *P < 0.001, one-way ANOVA with Bonferroni’s post hoc test. (**F**), ELISA analysis of IFN-β level in the supernatants of indicated primary neurons with the same treatment conditions. Mean ± SD; n =5; ^#^P < 0.05, *P < 0.01, one-way ANOVA with Bonferroni’s post hoc test.

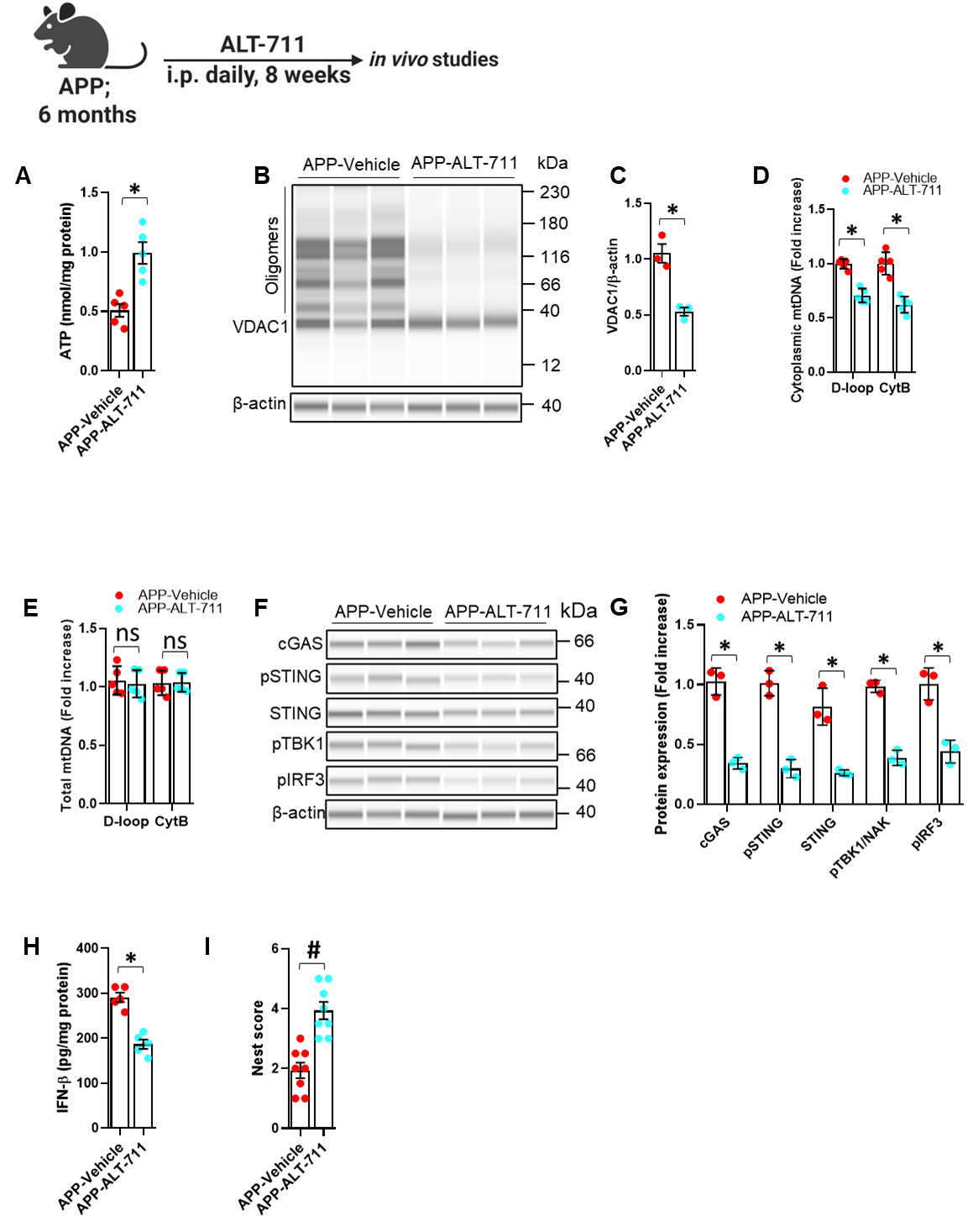

**Fig. S.11. ALT-711 mitigates mitochondrial dysfunction, mtDNA release, and cGAS-STING mediated neuroinflammation in APP mice.** (**A**), ATP levels were measured in cortical lysates from APP mice treated with or without ALT-711 (1 mg/kg BW) for 8 weeks, starting at 6 months of age. Mean ± SD; n = 5; *P < 0.01, two-tailed Student’s t-test. **(B, C)**, Immunoblotting analysis of the expression and quantification of VDAC1 oligomers relative to β-actin in indicated samples. Mean ± SD; n = 3; *P < 0.01, two-tailed Student’s *t*-test. (**D, E**), Quantification of cytoplasmic and total mitochondrial DNA (mtDNA) by qPCR in indicated APP mice samples. D-loop, and CytB are specific fragments or proteins encoded by the mitochondrial genome. Mean ± SD; n = 5; ^ns^P > 0.05, *P < 0.01, two-tailed Student’s *t*-test. **(F, G),** Immunoblot analysis for the expression and quantification of the expression of the cGAS, p-STING, STING, p-TBK1, and p-IRF3, relative to β-actin in the indicated samples. Mean ± SD; n = 3; *P < 0.001, two-tailed Student’s *t*-test. (**F**), ELISA analysis of IFN-β level in the indicated samples. Mean ± SD; n =5; *P < 0.01, two-tailed Student’s *t*-test. (**X**) Nest score based on nest building behavior. Nesting deficits indicated by incomplete nests and low nest score. Nesting ability indicated by complete nests, high nest score. Mean ± SD; n = 8; #P < 0.05, , two-tailed Student’s *t*-test.

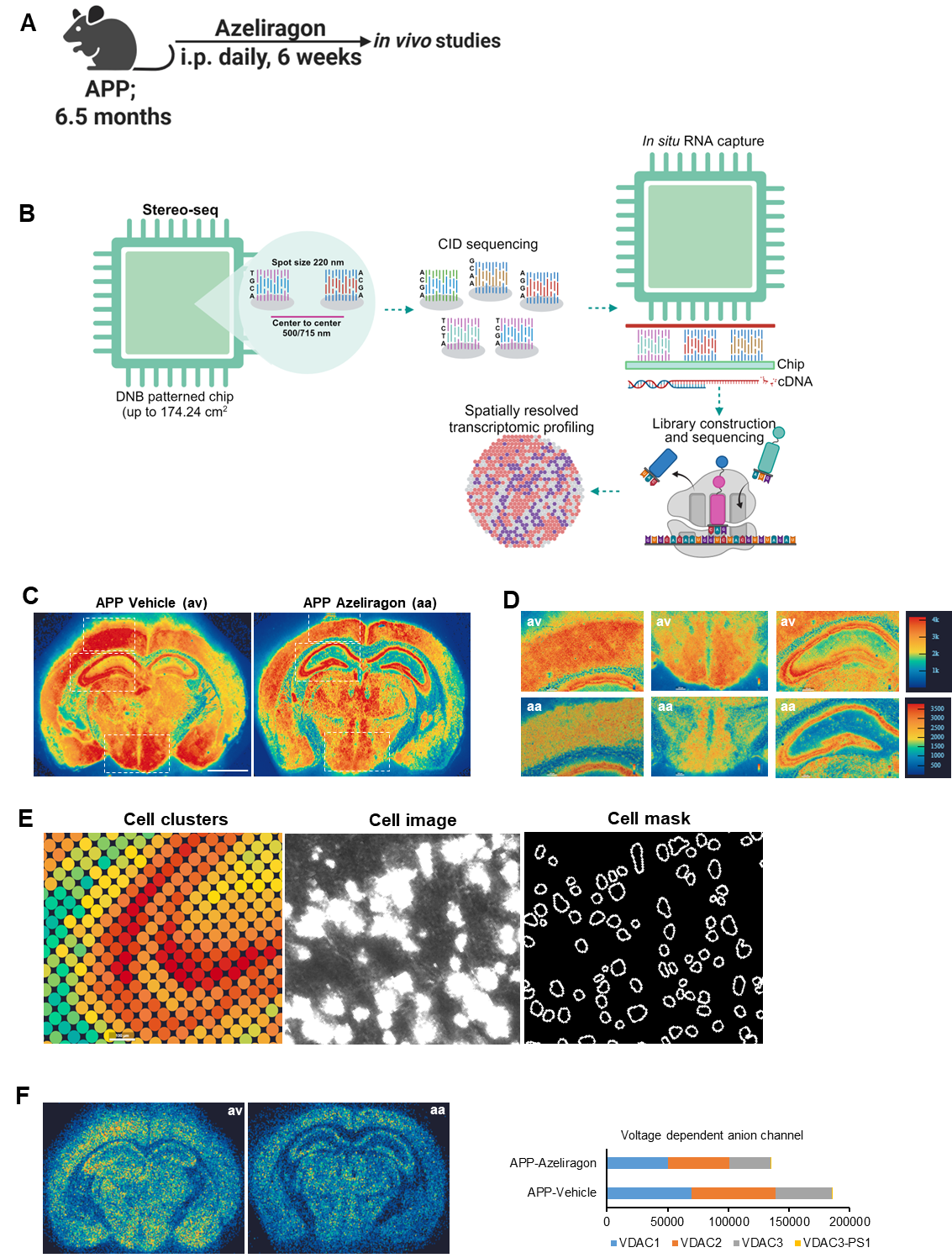

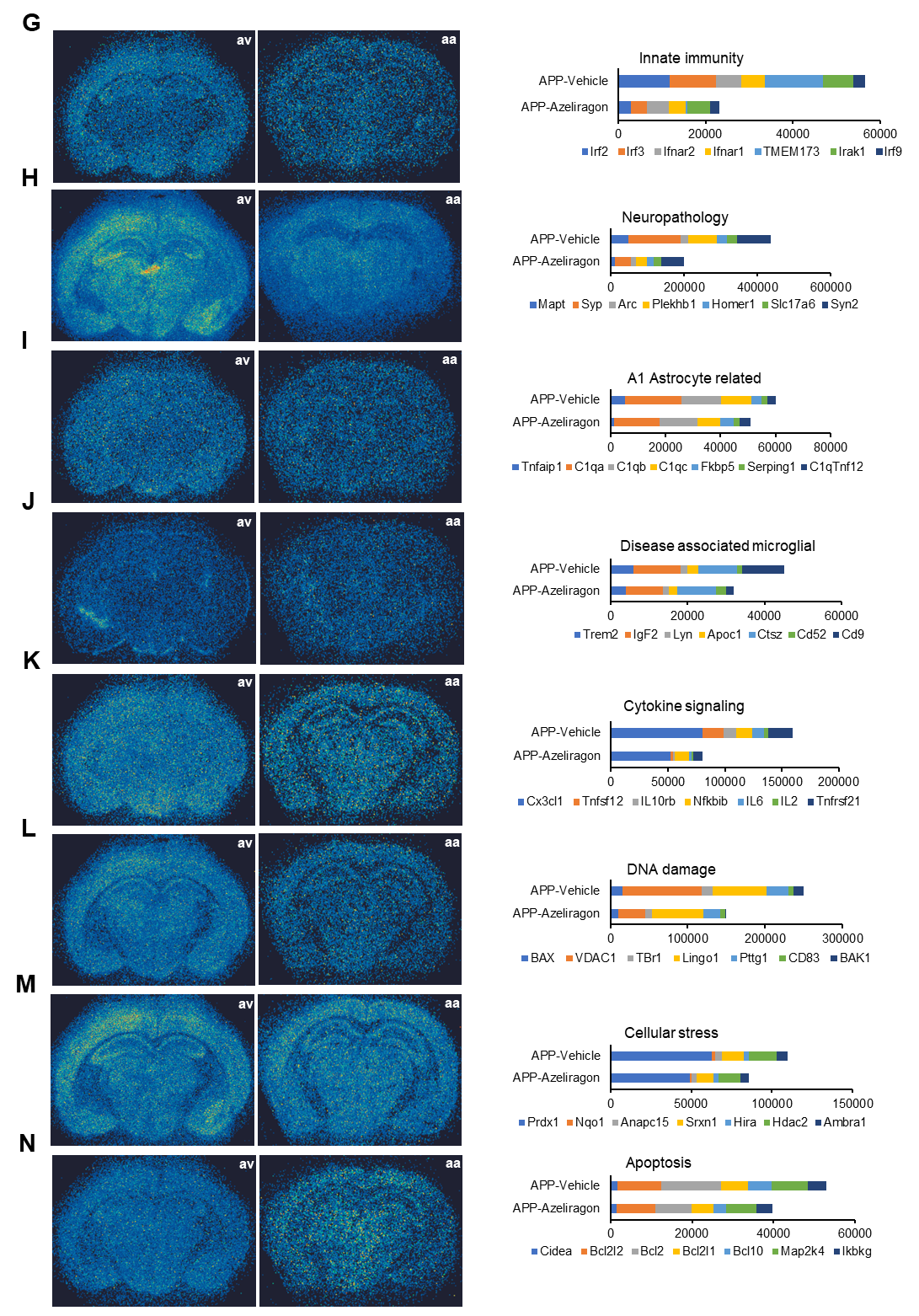

**Fig. S.12.** **Stereo-seq with spatial and cellular resolution in APP mice brains with or without RAGE inhibition.** (**A**) Schematic representation of vehicle or Azeliragon (a RAGE inhibitor) treatment on 6.5-month-old APP mice. (B) Stereo-seq pipeline: Design of the DNB patterned array chip, in situ sequencing to determine the spatial coordinates of uniquely barcoded oligonucleotides, preparation of capture probes by ligating the UMI-polyT containing oligonucleotides to each spot, in situ RNA capture from tissue, cDNA amplification, library construction, and sequencing and data analysis. (**C**) Spatial visualization of the detected cell clusters (scale bars, 100 µm), cell image, and cell mask in order to represent the DNA nanoball (DNB)-patterned array signal. (**D**) Stereo-seq: Enraged view of coronal section of indicated mice and the different regions (**E**) in order to show the changes in the MID count of genes. (**F-N**), Representative spatial visualizations of sections showing the spatial location and expression level (MID count difference) of genes related to components of voltage dependent anion channel (VDAC) (C), innate immunity (D), neuropathology (E), A1 astrocyte-related genes (F), disease-associated microglial (G), cytokine signaling (H), DNA damage (I), cellular stress (J) and apoptosis (K). MID count of genes is colored and labeled by cluster IDs with a 100% stacked column.

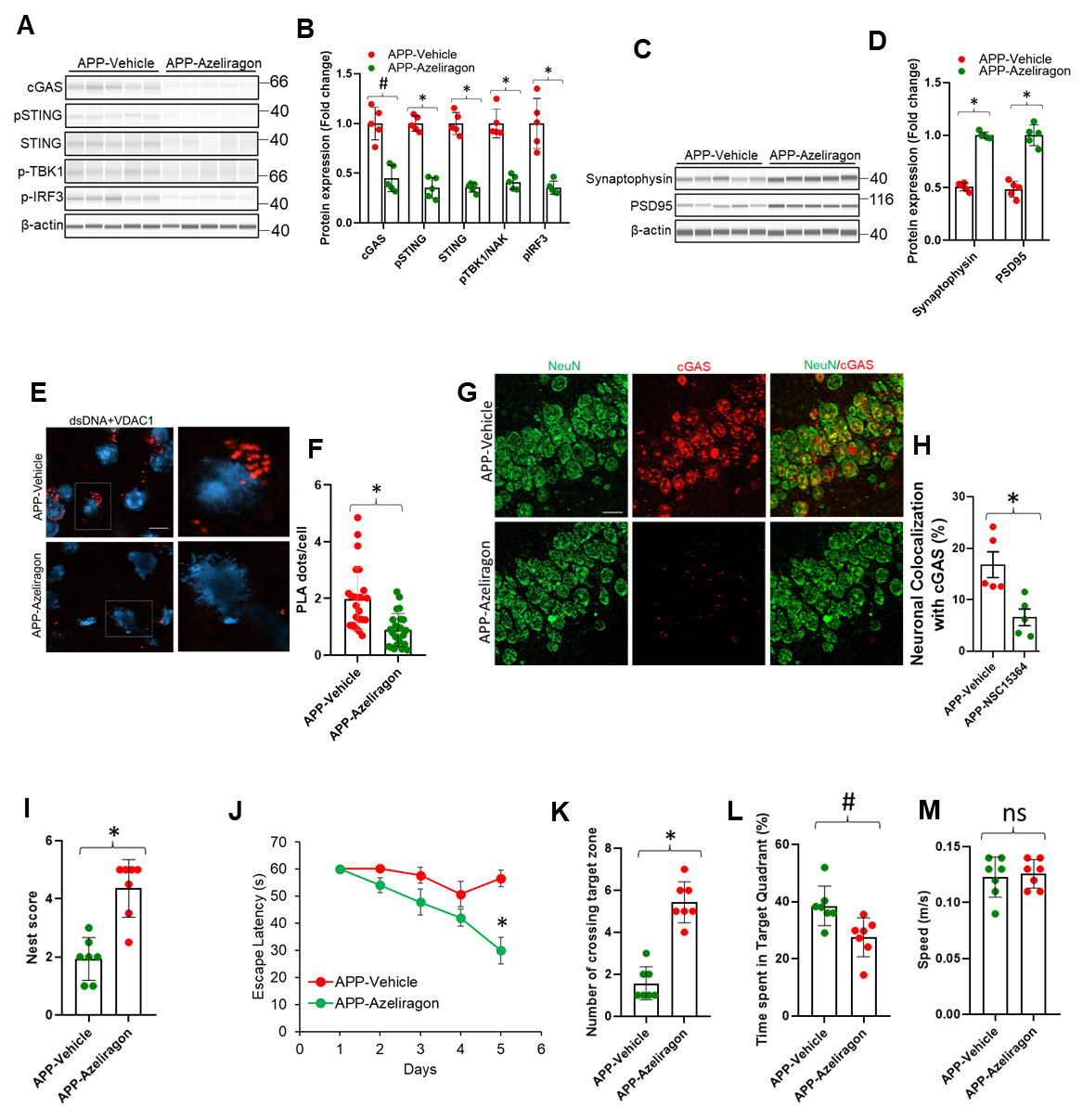

**Figure. S.13**. **RAGE inhibition rescues cGAS-STING activation, neurodegeneration, and memory impairment in APP mice.** (**A**) Immunoblotting analysis of the expression of the cGAS, p-STING, STING, p-TBK1, and p-IRF3, in six-week vehicle or Azeliragon treated APP mice (8-month-old). (**B**), Quantification of the expressions of cGAS, p-STING, STING, p-TBK1, and p-IRF3 relative to β-actin. Mean ± SD; n = 5; ^#^P < 0.05, *P < 0.01, one-way ANOVA with Bonferroni’s post hoc test. (**C**), Immunoblotting analysis of the expression of pre-synaptic marker protein synaptophysin and postsynaptic density protein 95 (PSD95) in cortical tissues (including hippocampus) (**D**), Quantification of protein levels of Synaptophysin or PSD95 relative to β-actin. Mean ± SD; n = 5; *P < 0.01, one-way ANOVA with Bonferroni’s post hoc test. (**E**), PLA with anti-dsDNA and anti-VDAC1 antibodies showing the association of VDAC1 protein with cytosolic or nuclear dsDNA in hippocampal CA1 region of the indicated mice. Scale bar, 10 μm. (**F**) Quantification of PLA red dots per cell. Five sections of each brain sample were stained and counted. Mean ± SD; n = 5; *P < 0.01, one-way ANOVA with Bonferroni’s post hoc test. (**G**), Immunostaining of cGAS with NeuN in Hippocampus of indicated mice. Scale bar, 10 μm. Representative images as shown were from the results of 5 mice in each group. (**H**) Percentage neuronal colocalization with cGAS. Mean ± SD; n = 5; *P < 0.01, one-way ANOVA with Bonferroni’s post hoc test. (**I**), Nest score based on nest building behavior. Nesting deficits indicated by incomplete nests and low nest score. Nesting ability indicated by complete nests, high nest score.   Mean ± SD; n = 7; *P < 0.01, one-way ANOVA with Bonferroni’s post hoc test. (**J-M**), Escape latency during MWM hidden platform task in indicated mice (J), mean number of crossings of the target during the probe test (K), time spent in the quadrant with the hidden platform (L), and average swim speed (M) of mice in MWM. Mean ± SD; n = 7; ^ns^P > 0.05, ^#^P < 0.05, *P < 0.01, one-way ANOVA with Bonferroni’s post hoc test.

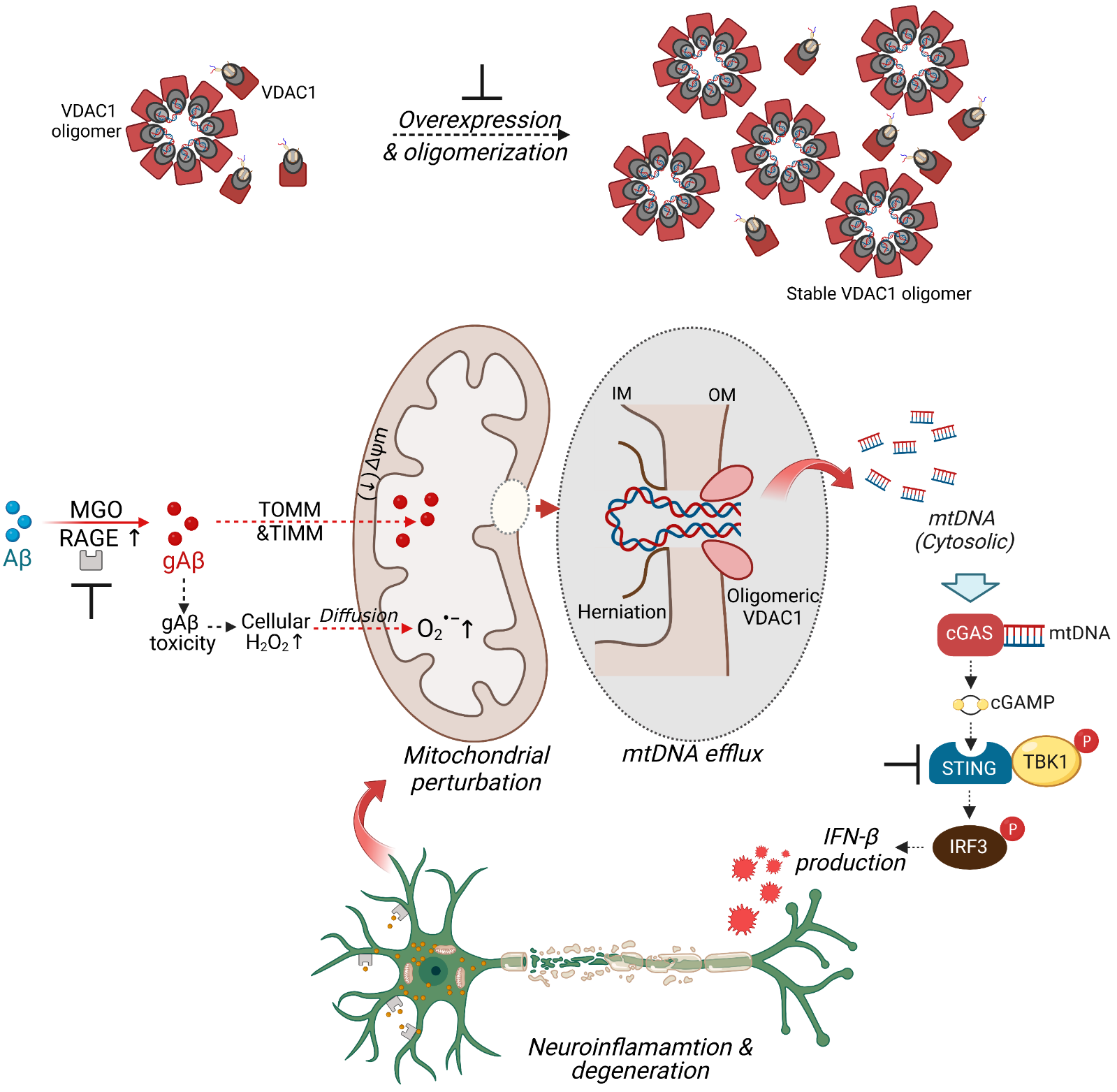

**Fig. S.14. Schematic representation of VDAC1 oligomerization and the cGAS-STING signaling pathway in neurons.** Aβ is a suitable substrate for highly reactive dicarbonyl (methylglyoxal; MGO)-induced glycation, forming glycated-Aβ (gAβ), which has a higher affinity with and uptake by the receptor of AGE (RAGE) in neurons. RAGE facilitates the mitochondrial superoxide generation and translocation of gAβ from the extracellular to the intracellular milieu and subsequently into mitochondria, aided by TOMM-TIMM (the translocase of the outer and inner mitochondrial membrane), thereby exacerbating gAβ-induced cytotoxicity through mitochondrial dysfunction. Increased glycation stress induces mitochondrial dysfunction by affecting the electron transport chain (ETC), decreasing mitochondrial membrane potential, and overexpressing and oligomerizing voltage-dependent anion-selective channel 1 (VDAC-1) accompanied by the N-terminal domain translocation into the large oligomer pore. VDAC1 oligomerization allowed the fragmented mitochondrial DNA (mtDNA) to reach the cytosol. Upon recognition of cytosolic mtDNA, cGAS catalyzes the formation of cGAMP and thereby activates STING. STING then activates and phosphorylates TBK1, which further phosphorylates IRF3. Phosphorylated IRF3 activates the expression of type-I IFNs in the nucleus, inducing neuronal inflammation, loss, and AD-like pathology.
